# Brown Adipose Tissue undergoes pathological perturbations and shapes C2C12 myoblast homeostasis in the SOD1-G93A mouse model of Amyotrophic Lateral Sclerosis

**DOI:** 10.1101/2024.07.29.605540

**Authors:** Marco Rosina, Silvia Scaricamazza, Flaminia Riggio, Gianmarco Fenili, Flavia Giannessi, Alessandro Matteocci, Valentina Nesci, Illari Salvatori, Daniela F. Angelini, Katia Aquilano, Valerio Chiurchiù, Daniele Lettieri Barbato, Nicola Biagio Mercuri, Cristiana Valle, Alberto Ferri

## Abstract

Amyotrophic lateral sclerosis (ALS) is a progressive neurodegenerative disease characterized by the selective loss of motor neurons. While the contribution of peripheral organs remains incompletely understood, recent evidence suggests that brown adipose tissue (BAT) and its secreted extracellular vesicles (EVs) could play a role in diseased context as ALS. In this study, we employed a multi-omics approach, including RNA sequencing (GEO identifier GSE273052) and proteomics (ProteomeXchange identifier PXD054147), to investigate the alterations in BAT and its EVs in the SOD1-G93A mouse model of ALS. Our results revealed significant changes in the proteomic and transcriptomic profiles of BAT from SOD1-G93A mice, highlighting ALS-related features such as mitochondrial dysfunction and impaired differentiation capacity. Specifically, primary brown adipocytes (PBAs) from SOD1-G93A mice exhibited differentiation impairment, respiratory defects, and alterations in mitochondrial dynamics. Furthermore, the BAT-derived EVs from SOD1-G93A mice displayed distinct changes in size distribution and cargo content, which negatively impacted the differentiation and homeostasis of C2C12 murine myoblasts, as well as induced atrophy in C2C12-derived myotubes. These findings suggest that BAT undergoes pathological perturbations in ALS, contributing to skeletal muscle degeneration through the secretion of dysfunctional EVs. This study provides novel insights into the role of BAT in ALS pathogenesis and highlights potential therapeutic targets for mitigating muscle wasting in ALS patients.

## 1. Introduction

Amyotrophic lateral sclerosis (ALS) is a neurodegenerative disease, characterized by upper and lower motor neuron degeneration, predominantly occurring in the adult age (50-70 years of age at onset) and leading to patients’ death within 2 to 5 years from diagnosis^1–3^. ALS is mainly characterized by progressive paralysis and atrophy of skeletal muscles, affecting also feeding behavior and the cardio-respiratory functionality. The etiology of ALS is very complex, due to the heterogeneity of the genetic background of patients with familiar ALS (approx. 10%) as well as the high occurrence of sporadic cases (approx. 90%)^3–6^. The function of genes involved in ALS pathology spans from antioxidant response (SOD1), proteostasis and autophagy (C9ORF72), and RNA metabolism (TARDBP, FUS), among an increasing number of newly discovered mutations (reviewed in ^7^)

The main metabolic feature of ALS is the higher energy expenditure, leading to an imbalance of the energetic status. ALS patients show lower body-mass index and reduced fat mass^8,9^. Moreover, retrospective studies have shown that weight loss can occur 10 years earlier than symptoms onset and weight loss entity positively correlates with faster disease progression and worse prognosis, suggesting that the metabolic defects could have both diagnostic and prognostic values^8,10,11^. Although defective food intake could be causative for weight loss during the clinical phase of the disease^12^, it does not justify the hypermetabolic state of ALS patients, as shown in indirect calorimetry assays, both in familial and sporadic cases^13,14^. Clinical biochemical characterization of ALS patients showed high incidence of hyperlipidemic status correlated with alteration of the cholesterol-lipoproteins ratios and overall defective lipid metabolism ^15^.

In this context, adipose tissue (AT) is the organ that systemically maintains the lipid energy homeostasis. AT accounts for the 10-30% of tissue body weight (depending on sex) and is responsible for the storage and utilization in all mammals. Two main types of anatomically and physiologically different AT are recognized: white AT (WAT) and brown AT (BAT). White adipocytes contain one large unilocular lipid droplet that occupies the largest part of the cytoplasm, leading the nucleus to the periphery of the cell body ^16^. Moreover, being mostly deputed to lipid storage, white adipocytes contain a very low number of mitochondria and are almost anabolic at rest ^17^. On the contrary, brown adipocytes contain many and smaller lipid droplets diffused into the cytoplasm and a very high number of mitochondria. Mitochondria are the cellular organelle providing the real physiological function of brown adipocytes ^18^. In fact, brown adipocytes display a high rate of fatty acid oxidation to maintain the highest level of mitochondria respiration and oxygen consumption. Differently from the canonical mitochondrial route, the proton-motive force is dissipated as heat by the uncoupling protein 1 (UCP1), reducing the net production of ATP by mitochondria ^19^. Consistently, ATP production in BAT is predominantly relying on glycolysis. BAT is essential for hibernating mammals, and in human pathophysiology correlates with healthier metabolic status and resistance to obesity and metabolic diseases since it provides removal of excess intake of dietary fatty acids (frequently occurring in western populations).

Given the metabolic role of BAT and the above reported metabolic defects in ALS patients, it is not surprising that the scientific community is considering this neglected tissue as a possible actor in ALS pathophysiology. Although the first reports about BAT modifications in the ALS context are available since the 2000s, the information and data provided are often confused and contradictory.

One BAT feature that is gaining even more attention by the scientific community is extracellular vesicles production that could mediate the cell-to-cell communication. BAT is an important source of circulating EVs, that can be altered in abundance and cargo according to physiological needs ^20–22^. Generally, in organs with high metabolic requirements, the mitochondria homeostasis is also maintained through active secretion of damaged mitochondrial particles ^23,24^.

In this study we show data partially filling the knowledge gap on the role of BAT in the pathogenesis of ALS, with particular attention to the alteration of the EVs production and their bioactive role on the homeostasis of skeletal muscle stem cells.

## 2. Material and methods

### 2.1. Animal model

All animal procedures were performed following the European guidelines for the use of animals in research (2010/63/EU) and the requirements of Italian laws (D.L. 26/2014); they were approved by the Italian Ministry of Health (protocol number 293/2021-PR).

Transgenic hemizygous SOD1-G93A male mice (B6.Cg-Tg [SOD1 G93A]1Gur/J) were obtained from The Jackson Laboratory (Bar Harbor, ME, United States of America; RRID:MGI:4835776) and then crossbred with C57BL/6 female mice to gain offspring. Generated animals were genotyped by PCR through hSOD1 oligos using interleukin-2 (IL-2) as a PCR internal control (sequences in Supplementary Table 1).

For this study male mice of 120 days post-partum (d.p.p.) of age were employed. Mice were euthanized through cervical dislocation.

### 2.2. RNA-preparation and RNA-sequencing

After euthanasia, mice were rinsed with 70% ethanol and BAT was exposed, removed with sterile scissors, and weighted. RNA was isolated with RNeasy Lipid Tissue Mini Kit (Qiagen, cat. #74804). Briefly, BAT tissue was homogenized in 1 ml of QIAzol Lysis Reagent. Phase separation was performed with addition of 200 μl of chloroform, vigorous shaking and centrifugation at 12000 ×g for 15 min at 4°C. Aqueous phase was extracted, and 1 vol of 70% ethanol was added to provide appropriate binding conditions. The sample is then applied to the RNeasy spin column and processed according to manufacturer’s instructions. Total RNA was eluted in 30 µl of RNase-free water. For real time qPCR experiments, RNA was prepared according to standard procedures using Trizol reagent (Thermo Fisher, 15596026). RNA concentration was assessed with NanoDrop Lite Spectrophotometer. The organic phase of RNA preparation was saved apart for subsequent isolation of total proteins (see above for details). Total RNA was quantified using the Qubit 4.0 fluorimetric Assay (Thermo Fisher Scientific). Libraries were prepared from 125 ng of total RNA using the NEGEDIA Digital mRNA-seq research grade sequencing service (Next Generation Diagnostic srl)^25^ which included library preparation, quality assessment and sequencing on a NovaSeq 6000 sequencing system using a single-end, 100 cycle strategy (Illumina Inc.). The raw data were analyzed by Next Generation Diagnostic srl proprietary NEGEDIA Digital mRNA-seq pipeline (v2.0) which involves a cleaning step by quality filtering and trimming, alignment to the reference genome and counting by gene^26–28^. The raw expression data were normalized, analyzed and visualized by Rosalind HyperScale architecture ^29^ (OnRamp BioInformatics, Inc.). Data are available at GEO identifier GSE273052

### 2.3. Total protein preparation and mass spectrometry analysis

The organic phase from RNA preparation was processed for total protein preparation, according to manufacturer’s instructions. Briefly, the interphase was removed and 0.3 ml of ethanol 100% were added, incubated at RT and centrifuged at 2000 ×g for 5 minutes at 4°C. the phenol-ethanol supernatant was transferred to a new tube and 1.5 ml isopropanol were added, incubated for 10 minutes at RT and centrifuged at 12000 ×g for 10 minutes at 4°C to pellet proteins. Protein pellets were washed for time with 0.3 M guanidinium hydrochloride solution in 95% ethanol, with 20 minutes of incubation at RT and centrifugation at 7500 ×g for 5 minutes at 4°C. The last wash was performed with 2 ml of 100% ethanol and the pellet was air-dried for 10 minutes. The pellet was resuspended in 0.2 ml of SDS 1% and heated for 15 minutes at 50°C to allow complete dissolution. Protein concentration was determined with Bradford assay. Protein leftover was processed for western blotting as described above. Instruments for LC-MS/MS analysis consisted of a NanoLC 1200 coupled via a nano-electrospray ionization source to the quadrupole-based Q Exactive HF benchtop mass spectrometer. Peptide separation was carried out according to their hydrophobicity on a PicoFrit column, 75 μm ID, 8 μm tip, 250 mm bed packed with Reprosil-PUR, C18-AQ, 1.9 um particle size, 120 Angstrom pore size (New Objective, Inc., cat. PF7508-250H363), using a binary buffer system consisting of solution A: 0.1% formic acid and B: 80% acetonitrile, 0.1% formic acid. Total flow rate: 300 nl/min. After sample loading, run start at 5% buffer B for 5 min, followed by a series of linear gradients, from 5% to 30% B in 90 min, then a 10 min step to reach 50% and a 5 min step to reach 95%. This last step was maintained for 10 min. MS spectra were acquired using 3E6 as an AGC target, a maximal injection time of 20 ms and a 120,000 resolution at 200 m/z. The mass spectrometer operated in a data dependent Top20 mode with subsequent acquisition of higher-energy collisional dissociation (HCD) fragmentation MS/MS spectra of the top 20 most intense peaks. Resolution for MS/MS spectra was set to 15,000 at 200 m/z, AGC target to 1E5, max injection time to 20 ms and the isolation window to 1.6 Th. The intensity threshold was set at 2.0E4 and Dynamic exclusion at 30 second. All acquired raw files were processed using MaxQuant (1.6.2.10) and the implemented Andromeda search engine. For protein assignment, spectra were correlated with the *Mus musculus* (v. 2021) including a list of common contaminants. Searches were performed with tryptic specifications and default settings for mass tolerances for MS and MS/MS spectra. The other parameters were set as follow:

- Fixed modifications: Propionamide(C)
- Variable modifications: Oxidation, Acetyl (N-term).
- Digestion: Tripsin
- Min. peptide length = 7
- Max. peptide mass = 470 Da
- False discovery rate for proteins and peptide-spectrum = 1%

For further analysis, the Perseus software (1.6.2.3) was used and first filtered for contaminants and reverse entries as well as proteins that were only identified by a modified peptide [First filter]. The LFQ Ratios were log-transformed, grouped and filtered for min.valid number (min. 3 in at least one group) [Second filter]. Missing values have been replaced by random numbers that are drawn from a normal distribution. Two-sample t-test analysis was performed with FDR = 0.05. Proteins with Log2 Difference ≥ ± 1 and q-value <0.05 were considered significantly enriched. Data are available at ProteomeXchange identifier PXD054147.

### 2.6. Isolation and characterization of BAT-extracellular vesicles

Extracellular vesicles was isolated as previously described^20^. Briefly, BAT was dissected, washed with sterile PBS and minced into small pieces in 0.5 ml of isolation medium (DMEM/F12, 1% Pen/Strep) and incubated in standard conditions for 16h. Supernatants were filtered through 30 μm nylon mesh and centrifuged at 600 ×g for 10 minutes at 4°C to remove debris. The resulting suspension was ultracentrifuged at 100000 ×g for 2h at 4°C in a swinging bucket rotor. For proteomics and western blotting, EVs pellets was resuspended in RIPA buffer and processed as described above. To assess both the concentration and size of the isolated vesicles, Nanoparticle-tracking analysis (NTA) was performed. All samples were quantified using a Nanosight NS300 (Malvern Panalytical, UK) equipped with a 532 nm laser. The acquisition was made using a camera level of 15 and involved capturing five 60-second videos for each sample, followed by analysis. The pellets were resuspended in pre-filtered PBS, and according to the manufacturer’s instructions, samples were appropriately diluted (range: 1:50 to 1:200) to achieve a reading-compatible concentration.

### 2.7. Integrated Functional Enrichment analysis of proteomics and RNA-Seq data

Upregulated genes from RNA-Seq and proteins from BAT-tissue proteomics were integrated into the Metascape online platform ^30^. Same procedure was performed for the integration of upregulated proteins from BAT-tissue and BAT-EVs proteomics. The resulting data were retrieved from the software and used for results description without further manipulation.

### 2.8. Isolation, culture, and treatment of brown primary pre-adipocytes

Isolation and culture of mouse primary brown pre-adipocytes (mPBAs) was performed according to Ngo et al, 2021 ^31^ with minor modifications. BAT was washed with sterile ice-cold PBS and minced into small pieces into 0.5 ml of collagenase solution (HBSS w/ Ca^2+^/Mg^2+^/glucose, 3.5% BSA, 1% Pen/Strep, Amphotericin B 500 ng/ml, Gentamycin 40 μg/ml). Tissue pieces were transferred into a sterile 15 ml tube containing 6.5 ml of the same solution (total 7 ml). Collagenase type I (Gibco, cat. # 17100017) and type II (Gibco, Cat # 17101015) were added at final concentration of 1 mg/ml. Tubes were placed under gentle agitation for 1h at 37°C, with vortexing every 15 minutes. Tissue homogenates were filtered with 100 μm nylon mesh and centrifuged at 250 ×g for 5 minutes at RT, inverted 10 times and centrifuged again. Supernatants were discarded and cell pellets were treated with 1 ml of red blood cell lysis buffer (NH_4_Cl 154 mM M, K_2_HPO_4_ 10 mM, EDTA 0.1 mM), incubated for 5 minutes at RT, resuspended into 13 ml of washing buffer (HBSS w/o Ca^2+^/Mg^2+^ w/ glucose, 3.5% BSA, 1% Pen/Strep, Amphotericin B 500 ng/ml, Gentamycin 40 μg/ml) and filtered through a 40 μm nylon mesh. Cells were centrifuged at 500 ×g for 5 minutes at RT and resuspended in amplification medium (Cytogrow, RESNOVA, Cat. # TGM-9001-B) and counted. 2×10^5^ cells were seeded on 100 mm Ø dish and cultured for 72-96h in standard conditions. Cultures were upscaled or downscaled according to the amount of cells isolated. After amplification, mPBAs were seeded for appropriate experiment in growth medium GM (DMEM/F12, 10% FBS, 1% Pen/Strep) at density of 1.4x10^5^ cells/cm^2^. After 3 days in GM, induction of brown adipogenic differentiation was performed for 48 hours in adipogenic induction medium AIM (GM + dexamethasone 1 μM, 3-isobutyl-methyl xanthine IBMX 0.5 mM, indomethacin 0.125 mM, rosiglitazone 1 μM, T3 1 nM and insulin 1 μg/ml). After the induction, differentiation was continued in adipogenic maintenance medium AMM (GM + rosiglitazone 1 μM, T3 1 nM and insulin 1 μg/ml) until day 8, with medium change every 2 days. For isoproterenol treatment, cells were stimulated for 16h with 1 μM isoproterenol.

### 2.9. Culture and treatment of C2C12 murine myoblasts

C2C12 murine myoblasts were purchased from ATCC (Ref. C2C12 CRL-1772™) and cultured in DMEM high glucose, supplemented with 10% FBS, 1 mM sodium pyruvate, 2 mM glutamine and 1% Pen/Strep. For differentiation, cells were seeded at 1×10^4^ cells/cm^2^ and allowed to reach confluence for 72 hours. Differentiation was induced with differentiation medium, consisting of DMEM high glucose, supplemented with 2% horse serum, 1 mM sodium pyruvate, 2 mM glutamine and 1% Pen/Strep. For treatments with BAT-EVs, vesicles were resuspended in sterile 1× PBS. Protein concentration was assessed with Bradford assay. Then, EVs were aliquoted and stored at -80. For growth curve experiment, cells were seeded at 5×10^3^ cells/cm^2^ and treated after over-night incubation with increasing concentration of BAT-EVs (0-0.5-1-2-4-8-16 μg/ml) for four time points (0-24-48-72 hours). For viability MTS experiment, cells were seeded at 4×10^4^ cells/cm^2^ and treated after over-night incubation with increasing concentration of BAT-EVs (0-0.5-1-2-4-8-16 μg/ml). for differentiation assay, cells were treated with 3.5 μg/ml of BAT-EVs and allowed to differentiate for 24 and 48 hours (for western blot) or up to 72 hours (for fusion index analysis). For the electron flow assay, cells were seeded at 4×10^4^ cells/cm^2^ and treated after over-night incubation with 3.5 μg/ml of BAT-EVs and analyzed after 24 hours. For atrophy experiments, myotubes were treated with 3.5 μg/ml of BAT-EVs and analyzed after 24 hours.

### 2.10. Cell lysis, protein extracts and western blotting analysis

Cells were washed 2 times with ice cold PBS and lysed in appropriate volume of RIPA buffer [Tris base 50 mM, 1% Triton X-100, 0.25% sodium deoxycholate, 0.1% SDS, NaCl 250 mM, 1 mM EDTA, 5 mM MgCl_2_, 1 mM sodium orthovanadate, 1 mM NaF, 1 mM phenylmethylsulfonyl fluoride, 1X Protease Inhibitor Cocktail (Sigma-Aldrich, #P8340), 1X Phosphatase Inhibitor Cocktail 2 (Sigma-Aldrich, #P5726), 1X Phosphatase Inhibitor Cocktail 3 (Sigma-Aldrich, #P0044)] and incubated in ice for 30 minutes. Lysates were sonicated using a probe sonicator and then centrifuged at 15,500 x g at 4 °C for 30 minutes. Protein quantitation was assessed with Bradford Bio-Rad Protein Assay Dye Reagent Concentrate (Bio-Rad, #5000006). Protein extracts were treated with 4X Laemmli buffer (0.25 M Tris base pH 6.8, 8% SDS, 40% glycerol, 20% 2-mercapto-ethanol, 4 mg/ml bromophenol blue), and heated at 56°C for 10 minutes, then stored at -20°C until used. 5-20 μg of proteins were loaded onto 4-20% SDS-polyacrylamide gel and run with tris-glycine-SDS buffer (0.025 M tris base, 0.192 M glycine, 0.01% SDS, pH 8.3). Gels were blotted onto nitrocellulose membranes (0.22 μm pore size) at 30V for 16h at 4°C in tris-glycine-ethanol buffer (0.025 M tris base, 0.192 M glycine, pH 8.3; 20% ethanol). Membranes were saturated with blocking buffer consisting of 5% non-fat dried milk in TBS-Tween buffer (0.02 M tris base, 0.15 M NaCl, 0.001 Tween-20). Primary antibodies were diluted appropriately in blocking buffer and incubated for 16h at 4°c under gentle agitation. Secondary antibodies were appropriately diluted in blocking buffer and incubated for 1h at RT under gentle agitation. Chemiluminescence was detected with Clarity™ Western ECL Substrate (Biorad) through iBright Imaging System (Thermo Fisher Scientific). Complete list of primary and secondary antibodies and dilutions is available in Supplementary Table 2. Densitometric analysis was performed with Fiji software ^32^.

### 2.11. cDNA synthesis and real time qPCR

cDNA synthesis was performed with ImProm-II™ Reverse Transcription System (Promega, cat. # A3800) according to manufacturer’s instructions. Real time qPCR analysis was performed with LightCycler 480 SYBR Green System (Roche, ETC). We used the “second derivative max” algorithm of the LightCycler software to calculate the crossing point (Cp) values. TATA box-binding protein as the housekeeping gene for normalization. Primer sequences are listed in Supplementary Table 3.

### 2.12. Adipocyte staining

Adipocyte staining was performed with Oil Red-O (ORO) as previously described^33–36^. ORO stock solution was prepared at 3 mg/ml in iso-propanol, stirred over night at 4°C and that filtered with standard filter paper. Cells were fixed with 4% paraformaldehyde solution, for 10 minutes at RT, washed 3 times with PBS, 1 time with ultrapure water and 1 time with 60% iso-propanol. Adipocytes were stained with appropriate volume of ORO working solution (OROstock:water 3:2, 0.22 μm filtered) for 20 minutes at RT under gentle agitation. After staining, nuclei were counterstained with DAPI following standard procedure. Images were acquired with ZOE Fluorescent Cell Imager (Biorad). ORO positive area and nuclei number was analyzed with a dedicate pipeline in Cell Profile software ^37^.

### 2.13. Immunofluorescence assay, fusion index analysis and myotube diameter

Cells were fixed with 4% PFA in PBS, for 10 minutes at RT and then washed with PBS. Permeabilization was performed with 0.5 % triton-X 100 in PBS for 5 minutes. Blocking was performed with 10% FBS and 0.1% Triton-X 100 in PBS for 1 hour under gentle agitation (5 rpm). Primary antibody anti myosin heavy chain (MF-20, DSHB) was diluted at 2 μg/ml in blocking solution and incubated for 2 hours at RT under gentle agitation (5 rpm). After 3 washes with 0.1% Triton-X 100 in PBS, secondary antibody (Goat anti-mouse IgG (H+L) cross-adsorbed secondary antibody, Alexa Fluor 488, Thermo Fisher Scientific, Cat. #A-11001) was diluted 1:500 in blocking solution and incubated for 1 hour at RT under gentle agitation. After 3 washing steps, nuclei were counterstained with Hoecsth 33342 (Invitrogen, Cat. no. H3570) diluted 1:5000 in PBS. Images were acquired with ZOE Fluorescent Cell Imager (Biorad) for analysis and with LSM 800 Confocal Laser Scanning Microscope (Zeiss) for representative images. Fusion index analysis was performed with a dedicated pipeline in Cell Profile Software ^37^. Myotube diameter analysis was performed with the Image J macro “Myotube width macro v.62” ^38^, with minor modification.

### 2.14. Mitochondrial morphological analysis

Primary brown pre-adipocytes were seeded at density of 1.4x10^4^ cells/cm^2^ (1/10 in respect to the differentiation protocol) and fixed with 4% PFA after over night incubation in standard culture condition. The mitochondrial network was labelled with a standard immunofluorescence protocol with anti-TOMM20 antibody (Supplementary Table 2) and goat anti-mouse Alexafluor 488 secondary antibody (Thermo Fisher Scientific, cat #A-11008). Images were acquired with Olympus BX51 Microscope. Mitochondrial network analysis was performed with the MiNA Plugin in the Image J software according to the standard analysis workflow indicated in the pubblication ^39^.

### 2.15. Bioenergetic analysis

Bioenergetic analysis was performed with the Seahorse XFe96 instrument (Agilent Technologies). mPBAs were seeded into Seahorse XF microplates (Agilent Technologies, Cat. #103794-100) at 15000 cells/well density in GM and cultured for 3 days in standard conditions. Differentiation protocol was performed as described above. Mitochondria-stress test assay was performed in Seahorse XF DMEM pH 7.4 medium (Agilent Technologies, Cat. #103575-100) supplemented with 1 mM sodium pyruvate, 10 mM D-glucose and 2 mM glutamine. Mitochondria were stressed with 1 μM oligomycin A, 1.5 μM FCCP and 1 μM rotenone/antimycin. Data were analyzed in the Seahorse Analyzer web platform (Agilent Technologies).

Electron flow assay on C2C12 myoblasts was performed in mitochondria assay solution (MAS), consisting of 220 mM mannitol, 70 mM sucrose, 10 mM KH_2_PO_4_, 5 mM MgC_l2_, 2 mM HEPES, 1 mM EGTA, 0.2% fatty acid-free BSA (Sigma Aldrich, cat. #A7030), 10 mM pyruvate, 1 mM malate, 4 mM ADP and 1 nM plasma membrane permeabilizer (Agilent Technologies, Cat #102504-100). Cells were incubated for 20 minutes in MAS, at 37°C in ambient air and analyzed immediately after. Complex I was inhibited with 2 μM rotenone, Complex II was stimulated with 10 mM succinate while Complex III was inhibited with 2 μM antimycin. Equilibration step was skipped. Measurements were performed 3 times with mix/wait/measures times of 30 sec/30 sec/2 min, for each injection step.

### 2.12. Mitochondrial potential and flow cytometry

Mitochondrial potential was measured by flow cytometry. Briefly, cells were washed in PBS and stained with MitoTracker Red (Molecular Probes, Eugene, OR) for 30 min in culture medium in standard culture conditions, at a final concentration of 500 nM according to manufacturer’s instructions. Mitochondrial fluorescence was analyzed with an CytoFLEX flow cytometer (Beckman Coulter). Excitation was achieved using the 640 nm laser. Gating parameters on the flow cytometer were established using a control sample and adjusting the voltages for the forward scatter, side scatter and MitoTracker Red laser. Data were analyzed by FlowJo software (Treestar, Ashland, OR, USA).

### 2.13. Statistical analysis

Statistical analysis for wet-lab experiments was performed in GraphPad Prism Software v10. Normality of datasets was assessed by D’Agostino-Pearson Test. Outliers were removed with ROUT method with Q=1%. Hetero/Homoscedasticity was assessed by the F test to compare variances. Comparison between two groups was performed with T-test, with appropriate correction for normality/non-normality and hetero/homoscedasticity as indicated in the relative figure legend. Comparison between more than two groups was performed by One-of Two-Way ANOVA test with multiple comparison correction when appropriate. For all comparisons, significant results were taken when p value < 0.05.

## 3. Results

### 3.1. Multi-omics analysis reveals ALS-related features in brown adipose tissue

Since the information about the role and the eventual modifications occurring at the level of BAT in the ALS contexts are often fragmented and contradictory, we decided to fill this gap with a complete characterization of BAT at the symptomatic stage of the disease, by a multi-omics approach involving both proteomics (ProteomeXchange identifier PXD054147) and RNA-Sequencing (GEO identifier GSE273052) analyses on BAT total extracts from wild type and SOD1-G93A mouse model at symptomatic stage of 120 d.p.p.. As expected, SOD1-G93A mice show a reduction in body weight, confirming that atrophic process, characteristic of this ALS model, has been through (**Fig. 1A**). Nevertheless, BAT did not show significant reduction in tissue mass neither at net weight nor normalized per body weight, suggesting that this tissue does not undergo excessive exhaustion at this stage (**Fig. 1B-C**).

**Figure 1:**
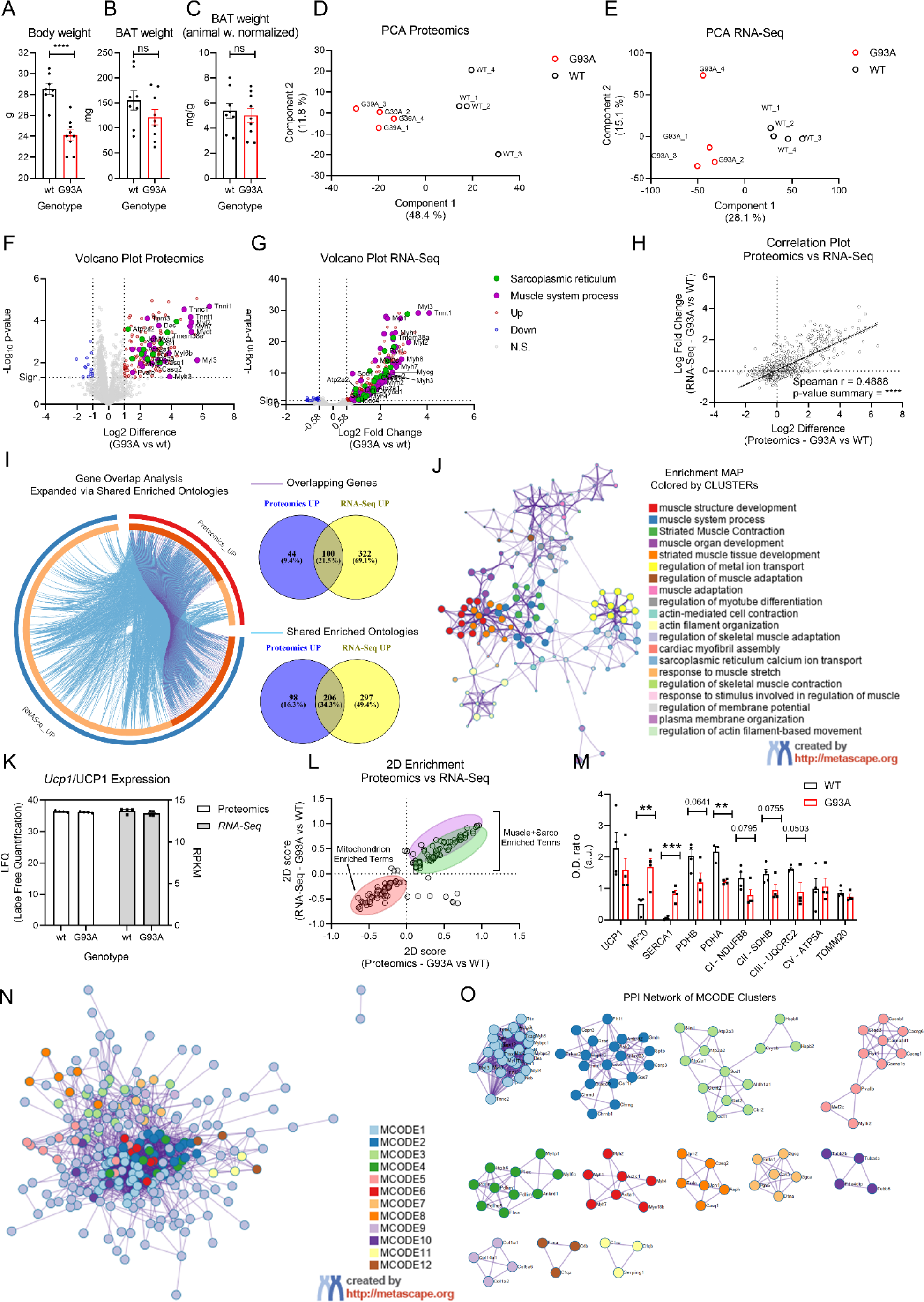
characterization of BAT from wild type and G93A mouse model. A-B-C) Bar plot representing mouse model weight, BAT raw and normalized weight. D-E) dispersion plot representing the principal component analysis of proteomics and RNA-Seq data. F-G) volcano plot of the proteomics and RNA-Seq data. H) dispersion plot representing the correlation analysis between proteomics and RNA-Seq datasets. I) chord diagram and Venn diagrams representing the genes and ontology overlap between proteomics and RNA-Seq up-regulated entities. J) enrichment map of commonly enriched ontology terms in proteomics an RNA-Seq up-regulated entities. Color coding identifies the different clusters of enriched terms. K) bar plot representing the raw quantification data of the UCP1/Ucp1 protein/gene in proteomics and RNA-Seq datasets, respectively. L) dispersion plot of the 2D enrichment analysis between proteomics and RNA-Seq data of BAT. M) bar plot representing the OD ration form densitometric analysis of western blot shown in Fig. S2 D-F. Significance was assessed by Student’s t-test. ** p<0.01; *** p<0.001. N) PPI network colored by MCODE clusters, relative to the Metascape integrated analysis of proteomics and RNA-Seq. O) PPI network of the separated MCODE clusters from panel N. Entities are reported as gene names.

To unveil molecular features characterizing the BAT from SOD1-G93A mice, we performed proteomics and RNA-Seq on total protein and RNA preparations. Correlation matrix showed high intragenotype similarity with proteomics Pearson coefficient 0.95 < r < 0.97 and RNA-Seq Pearson coefficient 0.97 < r < 0.98 (**Supp. Fig. 1A-B**). Principal component analysis (PCA) showed a sharp intergenotype segregation mainly across component 1, accounting for 48.4% and 28.1% in proteomics and RNA-Seq experiments, respectively (**Fig. 1D-E**). Differential expression analysis was performed separately for both datasets in the Perseus environment for proteomics ^40^ and in the Rosalind environment for RNA-Seq ^29^. Mass spectrometry allowed the identification of 3377 proteins, resulting in 2364 entities after filtering process. Among the 158 differentially expressed proteins (DEPs), 144 were overrepresented while 14 were underrepresented, considering q-value < 0.05 and |Log2 Difference| > 1 (**Fig. 1F** and **Supp. Fig. 1C**). RNA-Seq analysis allowed the identification of 437 differentially expressed genes (DEGs), consisting of 422 upregulated and 15 downregulated DEGs, considering adj. p-value < 0.05 and |fold change| > 1.5 (**Fig. 1G** e **Supp. Fig.** D). Since the two datasets showed positive correlation (**Fig. 1H**), enrichment and ontology analyses were performed in the Metascape on-line platform ^30^, considering the integration of overrepresented/upregulated entities from both datasets. The Gene Overlap Analysis Expanded via Shared Enriched Ontologies showed that the two datasets share 100 common entities and 206 enriched ontologies (**Fig. 1I**). Enriched ontology cluster network showed significant enrichment of terms mainly related to two macro-categories: acto-myosin complex and sarcoplasmic reticulum/calcium homeostasis (**Fig. 1J**, **Supp. Fig. 2A-C**). Interestingly, entities belonging to these categories are the ones driving the intergenotype segregation both in volcano-plots and in PCA analyses with particular focus on sarcolipin (*Sln*) gene being the predominant driver gene over component 1 in the RNA-Seq experiment (**Supp. Fig. 1E-F**). Among the mostly dysregulated entities, increased expression of myosin heavy/light chain isoforms and SERCA pumps predominantly characterized BAT from SOD1-G93A mouse model (**Supp. Fig. 1G-H**) suggesting that this tissue undergoes a molecular rearrangement imputable to the activation of alternative/non-canonical thermogenic mechanisms. This hypothesis is corroborated by the absence of modulation of Ucp1/UCP1 gene/protein expression in SOD1-G93A BAT (**Fig. 1K-L**). To deeper investigate the overall alterations in the molecular environment of BAT in the ALS context, we performed 2D-Enrichment analysis between total proteome and transcriptome in the Perseus platform^40^. Results confirmed the positive enrichment of Actomyosin-and Sarcoplasmic Reticulum-related terms, while also showing depletion of Mitochondria-related ones (**Fig. 1L** and **Supp. Fig. 1I**). Although mitochondria genes and proteins did not result from the previous statistical analyses, this information was confirmed by 2D Fisher test with Benjiamini-Hoechberg FDR cut-off < 0.02. Western blot analysis on total protein extracts from BAT confirmed the increase in abundance of sarcomeric myosin heavy chain (MHC) and SERCA1 ATPase (**Fig. 1M**, **Supp. Fig. 1D**) and corroborates the information related to decreased abundance of mitochondrial proteins, relatively to the inner matrix metabolic enzymes, pyruvate dehydrogenase subunit alpha (PDHA), pyruvate dehydrogenase subunit beta (PDHB), pyruvate carboxylase beta (PCB) (**Fig. 1M**, **Supp. Fig. 1E**) and electron transport chain complexes (**Fig. 1M**, **Supp. Fig. 1F**).

To gain further insights into the dynamics occurring at the molecular level between the dysregulated entities resulted from proteomics and RNA-Seq analyses, we adopted a network approach. Protein-protein interaction network analysis showed high interconnection between entities from both proteomics and RNA-Seq (**Supp. Fig. 2G**). MCODE clustering analysis recognized 12 different clusters in the PPI network (**Fig. 1** N-O, **Table 1**). MCODE1, MCODE2, MCODE4, MCODE6 and MCODE7 represented ontology terms related to the acto-myosin complex and muscle-related terms suggesting the activation of alternative thermogenic program through myosin-dependent futile ATP consumption. MCODE 3, MCODE5, and MCODE8 showed enrichment for pathways related to the intracellular calcium homeostasis, consisting in overexpression of SERCA pumps (Atp2a1, Atp2a2, Atp2a3), mitochondrial casein kinase (Ckmt2), mitochondrial and cytosolic transamination (Got1, Got2) and antioxidant response to lipid peroxidation (Aldh1a1), calcium voltage-gated channels (CACNs), ryanodine receptor (Ryr1), parvalbumin (Pvalb) and calsequestrins (Casq1, Casq2). These data suggest that calcium homeostasis is perturbed in BAT from SOD1-G93A mouse model and could be pivotal for the activation of ATP-consuming pathways contributing to alternative thermogenesis. Interestingly, MCODE9, MCODE10, MCODE11 and MCODE 12 report enrichment for other altered pathways such as collagen deposition, intracellular transport and the complement cascade, which could suggest dysregulation in parallel cellular functions that need further investigation.

**Table 1:** MCODE clustering details. Table reporting the list of MCODE cluster recognized by the integrated Metascape analysis of proteomics and RNA-Seq data showed in Fig. 1 N-O. For each cluster, the top 3 enriched terms are reported, relatively to the lowest p-value.

|  | Annotation |  |  |
| --- | --- | --- | --- |
| Network | Term ID | Term Description | p-value<br>(-Log10) |
| MCODE1 | R-MMU-390522 | Striated Muscle Contraction | -69.5 |
|  | R-MMU-397014 | Muscle contraction | -48.7 |
|  | GO:0006936 | muscle contraction | -48.6 |
| MCODE2 | GO:0061061 | muscle structure development | -11 |
|  | GO:0042692 | muscle cell differentiation | -8 |
|  | GO:0030036 | actin cytoskeleton organization | -6.6 |
| MCODE3 | R-MMU-418359 | Reduction of cytosolic Ca<sup>++</sup> levels | -7.2 |
|  | GO:0090075 | relaxation of muscle | -6.7 |
|  | R-MMU-418360 | Platelet calcium homeostasis | -6.3 |
| MCODE4 | GO:0061061 | muscle structure development | -13.2 |
|  | GO:0030029 | actin filament-based process | -8.9 |
|  | GO:0030036 | actin cytoskeleton organization | -7.3 |
| MCODE5 | mmu04921 | Oxytocin signaling pathway - Mus musculus (house mouse) | -15.6 |
|  | GO:1902514 | regulation of calcium ion transmembrane transport via high voltage-gated calcium channel | -10.1 |
|  | GO:0070588 | calcium ion transmembrane transport | -10 |
| **MCODE6** | mmu04814 | Motor proteins - Mus musculus (house mouse) | -14.3 |
|  | GO:0006936 | muscle contraction | -8.8 |
|  | GO:0003012 | muscle system process | -8.2 |
| **MCODE7** | CORUM:349 | Sarcoglycan-sarcospan-syntrophin-dystrobrevin complex | -12.9 |
|  | mmu05412 | Arrhythmogenic right ventricular cardiomyopathy - Mus musculus (house mouse) | -11.3 |
|  | mmu05416 | Viral myocarditis - Mus musculus (house mouse) | -11 |
| **MCODE8** | GO:1901019 | regulation of calcium ion transmembrane transporter activity | -11.8 |
|  | GO:0051279 | regulation of release of sequestered calcium ion into cytosol | -11.4 |
|  | GO:2001257 | regulation of cation channel activity | -10.8 |
| **MCODE9** | R-MMU-8948216 | Collagen chain trimerization | -11.2 |
|  | R-MMU-2022090 | Assembly of collagen fibrils and other multimeric structures | -10.7 |
|  | R-MMU-1650814 | Collagen biosynthesis and modifying enzymes | -10.2 |
| **MCODE10** | R-MMU-190840 | Microtubule-dependent trafficking of connexons from Golgi to the plasma membrane | -8.8 |
|  | R-MMU-190872 | Transport of connexons to the plasma membrane | -8.8 |
|  | R-MMU-9668328 | Sealing of the nuclear envelope (NE) by ESCRT-III | -8.2 |
| **MCODE11** | R-MMU-977606 | Regulation of Complement cascade | -8.3 |
|  | GO:0006958 | complement activation, classical pathway | -8.3 |
|  | GO:0002455 | humoral immune response mediated by circulating immunoglobulin | -8.1 |
| **MCODE12** | R-MMU-166663 | Initial triggering of complement | -9.1 |
|  | R-MMU-166658 | Complement cascade | -8 |
|  | GO:0006956 | complement activation | -7.8 |

**Supplementary Figure 1:**
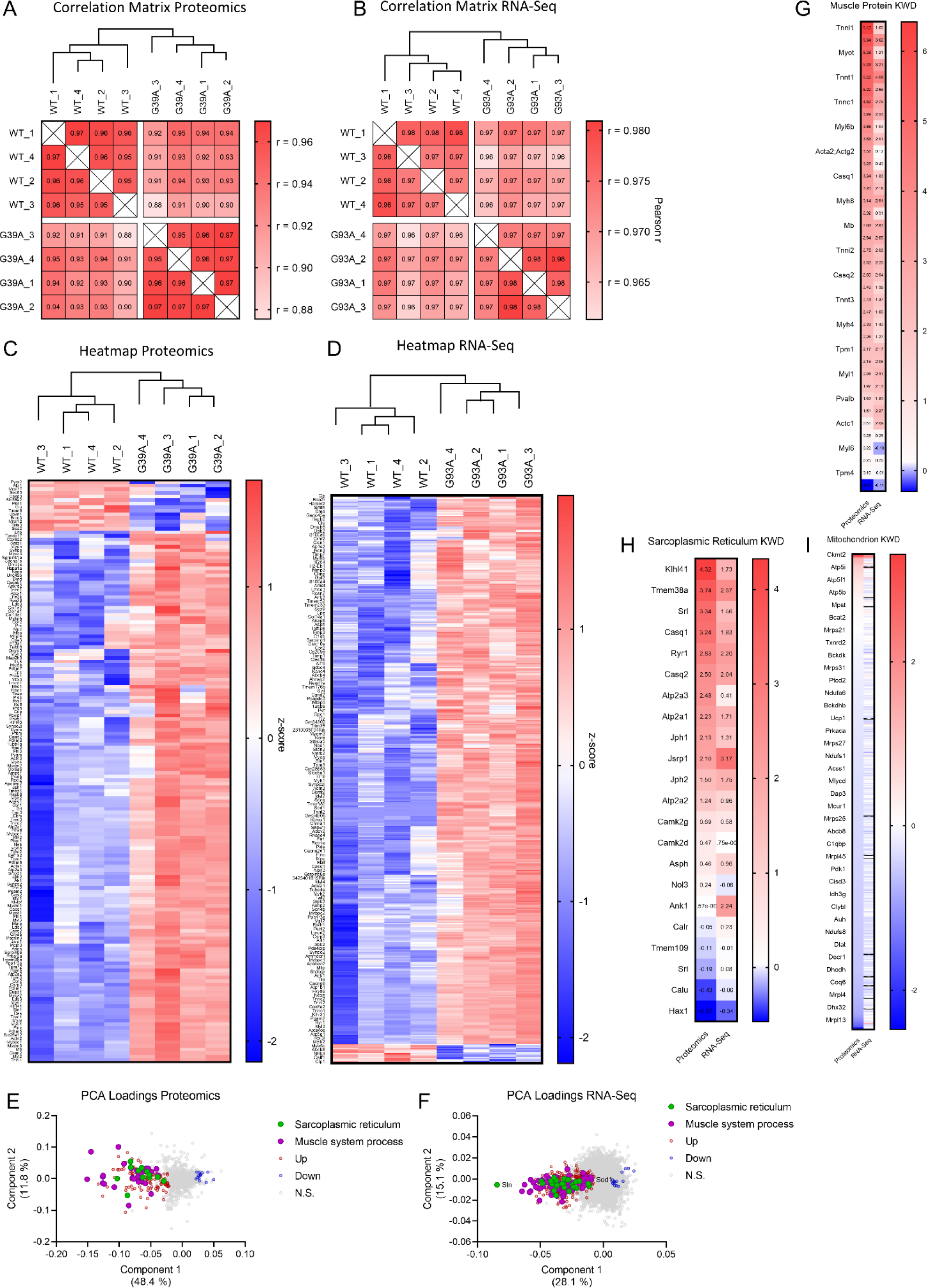
Proteomics and RNA-Seq supplementary information. A-B) heatmaps representing the Pearson correlation r values for dataset in proteomics and RNA-Seq analyses. C-D) heatmap representing the significantly regulated entities (proteins and genes, respectively) from proteomics and RNA-Seq analyses. Values are normalized in z-scoring and plotted as red (for up-regulated) and blue (for down-regulated).E-F) dispersion plots representing the distribution of entities (proteins and genes, respectively) according to Components 1 and Component 2 in the PCA analysis of Figure 1 D-E. G-H-I) heatmap representing the z-scored expression values of genes belonging to the relative Gene Ontology category. Values are plotted as red (for up-regulated) and blue (for down-regulated).

**Supplementary Figure 2:**
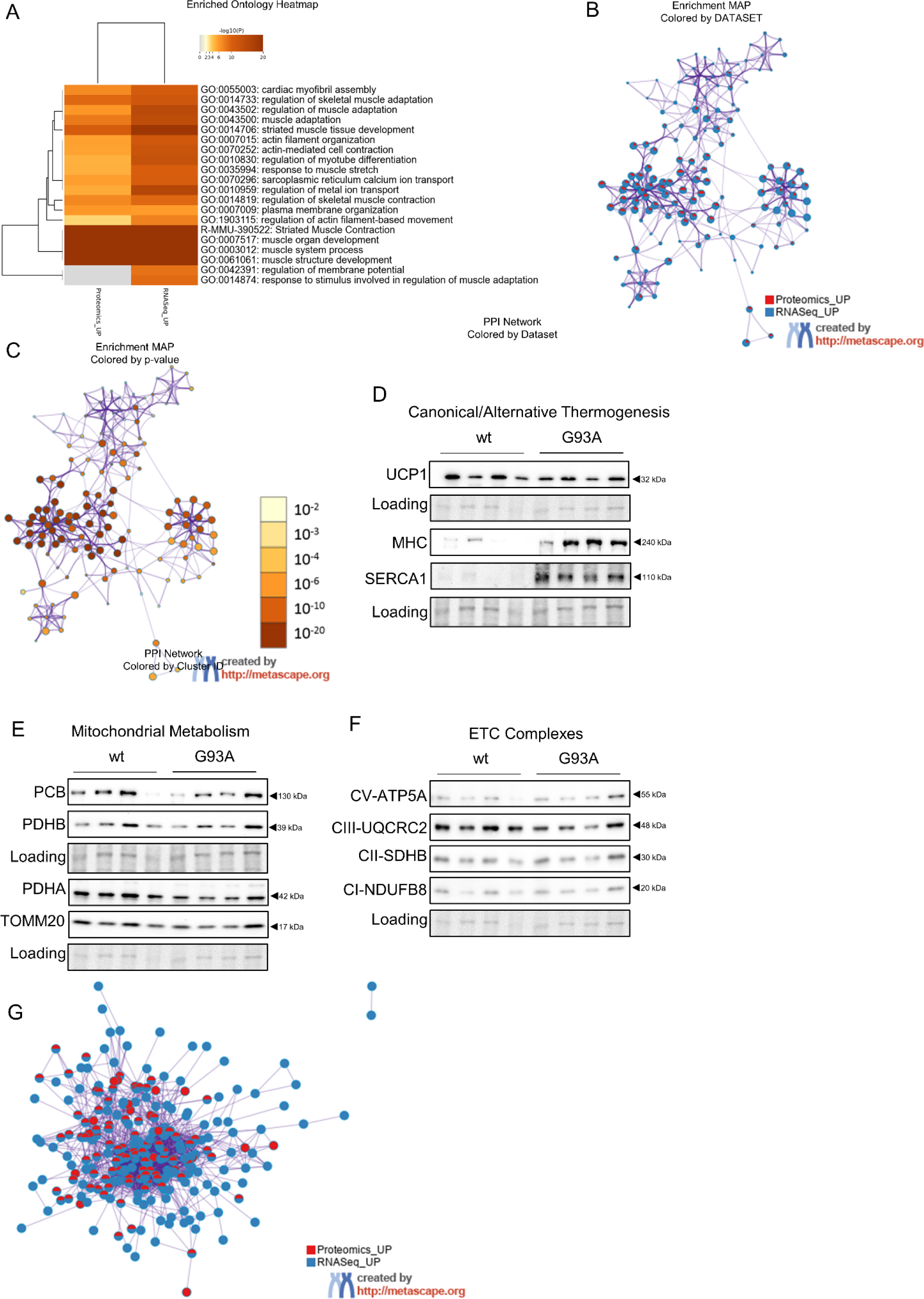
Gene Ontology supplementary information and Clustering. A) heatmap representing the enriched gene ontology -Log10 p-values in the Metascape analysis of integrated proteomics and RNA-Seq analyses upregulated entities. B) enrichment map colored by dataset (proteomics upregulated in red and RNA-Seq upregulated in blue), relative to Fig. 1J. C) enrichment map colored by p-value relative to Fig. 1J. D) representative western blot analysis of total BAT extracts for canonical (UCP1) and alternative (MHC, SERCA1) thermogenic markers. Ponceau S staining was taken as loading control. E) representative western blot analysis of total BAT extracts for mitochondrial proteins (PCB, PDHA, PDHB and TOMM20). Ponceau S staining was taken as loading control. F) representative western blot analysis of total BAT extracts for mitochondrial ETC complexes (NDUFB8, SDHB, UQCRC2 and ATP5A). Ponceau S staining was taken as loading control. G) protein-protein interaction (PPI) network, colored by dataset (proteomics upregulated in red and RNA-Seq upregulated in blue), relative to the Metascape integrated analysis.

### 3.2. Primary brown adipocytes from SOD1-G93A mice display differentiation impairment and respiratory defects

Multi-omics analysis highlighted several alterations in BAT from symptomatic SOD1-G93A mouse model, mainly consisting of activation of alternative thermogenic pathways and impairment of the mitochondrial compartment. To better characterize such defects, we performed experiments aimed analyzing the phenotypical demeanor and the metabolic functionality of *ex vivo* cultured murine primary brown adipocytes (mPBAs). mPBAs from SOD1-G93A mice showed reduced adipogenic index, expressed as Oil Red-O positive area normalized per total number of cells in fluorescence microscopy imaging (**Fig. 2 A-B**). Moreover, morphological screening of adipocytes through automated image analysis showed reduced adipocyte area (**Fig. 2 C-D**). In particular, mPBAs showed reduced differentiation and almost absence of adipocytes with area greater than 400 μm^2^ (**Fig. 2 D**). To obtain further information on the molecular signaling controlling adipogenic differentiation we performed western blot analysis throughout the differentiation process. After 48 hours from induction of differentiation with AIM, the transcriptional cascade controlled by PPAR γ1/γ2 and CEBPα was downregulated (Fig. 2 E,H). In particular, PPAR γ2 showed the lowest expression, compared to the γ1 isoform, suggesting a differential regulation between lipogenesis and mitochondrial metabolism. At terminal differentiation, after other 6 days in AMM, we dissected different aspects of the adipogenic differentiation. Firstly, we looked at late lipogenic markers as fatty acid binding protein 4 (FABP4), fatty acid synthase (FASN) and phosphorylated acetyl CoA-carboxylase (pACC Ser_79_). Notably, ACC phosphorylation on Ser_79_ resulted to be downregulated, suggesting an impaired lipid anabolic process (**Fig. 2 F, H**). We next analyzed the expression of mitochondrial metabolic markers (**Fig. 2 G-H**). As showed in the proteomic and transcriptomic profiling, UCP1 protein levels do not change consistently in SOD1-G93A mPBAs. Similarly, mitochondrial mass marker TOMM20 remains stable, suggesting that variation in mitochondrial proteins and genes highlighted by the multi-omics characterization does not involve the total mitochondrial network, but relies on the downregulation of mitochondrial metabolic proteins, as for the case of PDHA, PDHB and PCB. Moreover, antioxidant response, mediated by SOD2 protein does not seem to be activated, suggesting absence of increased redox flux in SOD1-G93A mPBAs. Overall, phenotypical characterization clearly shows adipogenic, lipogenic and metabolic defects in mPBAs from SOD1-G93A mouse model. Since multi-omics analysis and western blot showed alteration at the mitochondrial compartment, we next performed Extracellular Flux analysis through Seahorse Technology™ in order to assess the mitochondrial functionality (**Fig. 2 I**). Results show a drastic reduction in mitochondrial respiration (**Fig. 2 J**), with particular modulation of Basal and Maximal Respiration, Spare Capacity and Proton Leak (**Fig. 2K**), confirming an impairment of mitochondrial functionality (**Fig. 1 L-M**). Modulation in Basal and Maximal Respiration prompted us to assess the abundance of mitochondrial ETC complex by western blot. Interestingly, Complex I subunit NDUFB8 is downregulated in SOD1-G93A mPBAs (**Fig. 2 L**). This result could be correlated with respiration alterations, as Complex I downregulation could be responsible for a lower reducing-equivalent flux through the electron transport chain. This could be at the basis of also lower Spare Capacity and Proton Leak, as confirmed by the analysis of MitoTracker Red staining by flow-cytometry, showing reduced mitochondrial potential (**Fig. 2 M**). Overall, the results highlight that mPBAs from SOD1-G93A mice have differentiation impairment, probably due also to a lower mitochondrial performance unable to support a proper adipogenic differentiation. These defects could also affect cellular response to external stimuli.

**Figure 2:**
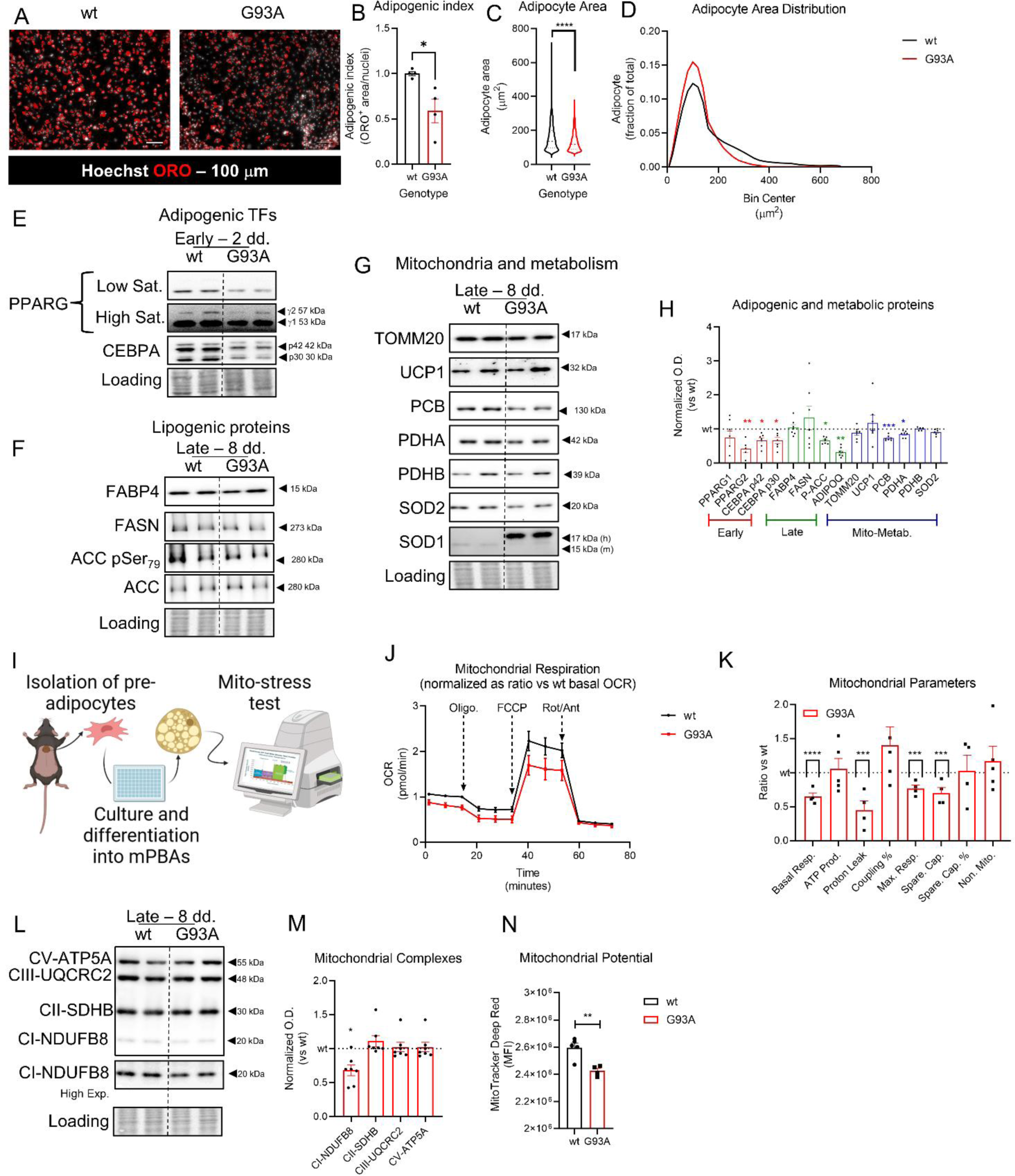
characterization of mPBAs from SOD1-G93A mouse model. A) representative fluorescent micrographs of mPBAs from wild type and SOD1-G93A mouse models. lipid content was labelled with oil-red O and nuclei were counterstained with Hoechst 33342. Scale bar = 100 μm. B) bar plot representing the quantitation of the adipogenic index relative to experiment in panel A. data were obtained as ORO-positive area normalized per nuclei and expressed as mean ± SEM. Statistical significance was assessed through Welch’s test. * p<0.05. n = 4 C) violin plot representing the quantitation of adipocyte area from images panel A. data were obtained through automatic image analysis and expressed as μm^2^. Dashed line represents the median value. Dotted lines represent the 1^st^ and the 3^rd^ quartiles. Statistical significance was assessed through Kolmogorov-Smirnoff test. **** p<0.0001. D) distribution plot of adipocytes area from images in panel A. data were obtained as for panel C and represented as frequency distribution. E) representative western blot analysis of adipogenic transcription factors PPARG and CEBPA analyzed at 48 hours after adipogenic induction. PPARG is shown at low and high saturation level to allow the appreciation of both the bands of isoform 1 and 2. Ponceau red staining was used as loading control. F) representative western blot of late lipogenic markers FABP4, FASN, ACC and ACC p-Ser79, analyzed at 8 days of differentiation. Ponceau red staining was used as loading control. G) representative western blot of late mitochondrial and metabolic markers TOMM20, UCP1, PCB, PDHA, PDHB, SOD2 analyzed at 8 days of differentiation. SOD1 antibody was used as internal control for human SOD1 overexpression in SOD1-G93A mouse model. Ponceau red staining was used as loading control. H) bar plot representing the densitometric quantitation of western blot experiments in panels E-G. data were obtained by normalization versus loading control and expressed as ration versus the mean of wild type samples. Statistical significance was assessed through Student’s t-test. * p<0.05; ** p<0.01; **** p<0.0001. nwt = 4; nG93A = 7. I) cartoon representing the experimental model for Seahorse experiment. Pre-adipocyes were isolated from BAT and expanded in Cytogrow. Thereafter, cells were seeded onto Seahorse microplates and let to differentiate into mPBAs for 8 days. Mito-stress test was conducted with Oligomycin 1 μM, FCCP 1.5 μM and Rotenone/Antimycin A 1/1 μM. artwork was mad with Biorender www.biorender.com J) graph representing the oxygen consumption rate (OCR) of mPBAs from wild type and SOD1-G93A mice during the mito-stress test, as described in panel I. data are reported as time on the x axis, measured in minutes, and OCR on the y axis, normalized versus the baseline value of wild type samples. K) bar-plot representing the mitochondrial respiratory parameters from Seahorse experiments. Data were obtained through the Seahorse analytics platform www.seahorseanalytics.agilent.com. Data are reported as mean ± SEM and normalized versus the mean on wild type samples. Statistical significance was assessed through the Student’s t-test. *** p<0.001; **** p<0.0001. n=5. L) representative western blot of mitochondrial ETC complexes NDUFB8, SDHB, UQCRC2 and ATP5A. NDUFB8 is shown also at higher exposure time. M) bar plot representing the densitometric quantitation of western blot in panel L. data are expressed as mean ± SEM and normalized versus the mean value of wild type samples. Statistical significance was assessed through Student’s t-test. * p<0.05 Wild type is shown as reference dotted line at y=1. n_wt_ = 4; n_G93A_ = 7. N) bar plot representing the quantitation of mitochondrial transmembrane potential through mitotracker deep red staining in flow-cytometry. Data are reported as mean fluorescent intensity ± SEM. Statistical significance was assessed through Student’s t-test. ** p<0.01. n = 5.

### 3.3. mPBAs from SOD1-G93A mice have alterations of mitochondrial dynamics and response to pro-lipolytic stimuli

The data shown in Figure 2 highlighted the presence of mitochondrial damage, impacting on both cellular differentiation and metabolism. To get deeper information about the status of the mitochondrial network, we performed confocal microscopy and MiNA analysis on mPB pre-adipocytes (Fig. 3A). Analysis of the skeletonized network showed reduction of mitochondrial individuals, as either puncta or rods (Fig. 3B). Moreover, a reduction of the number of networks was highlighted by the analysis, although the distribution of the entity type was similar between wild type and SOD1-G93A pre-adipocytes (Fig. 3 B-C). Notwithstanding, mitochondrial footprint analysis does not show a reduction in total mitochondrial area, suggesting that such alterations in network and entities abundance is not correlated to a reduction in mitochondrial mass, as also suggested by western blot analysis of TOMM20 on differentiated mPBAs (Fig. 2G). Interestingly, mitochondrial network analysis showed an increased network size, correlating with increased number of junctions and branch length (Fig. 3E). This result prompted us to hypothesize that mitochondrial network undergoes an increased fusion process in the G93A context. To test this hypothesis, we assessed the expression of DRP1 and OPA1 proteins in cell lysates of mPBAs at terminal differentiation (Fig. 3 F-G). Results show that DRP-1 phosphorylation on Ser-616 is reduced in mPBAs from SOD1-G93A mouse model. In parallel no consistent changes in OPA1 levels was observed. We hypothesize that such changes in the mitochondrial dynamics could also impact on the cellular competence to respond to pro-lipolytic stimuli. To test this hypothesis, we started with the analysis of the lipolytic signaling cascade into BAT tissue lysates. To assess the activation of the phosphorylation cascade driven by protein kinase A (PKA), under control of the adrenergic beta-3 receptor, we used an antibody directed against the PKA target epitope (RRXS*/T*). In addition, we analyzed the phosphorylation of the hormone-sensitive lipase HSL, a direct target of PKA on the activating Ser_563_. Results show that BAT from SOD1-G93A mouse model has an increased amount of PKA phospho-substrates, while presenting a significantly decreased phosphorylation of HSL Ser_563_ (**Fig. 3H-I**). In parallel, data do not show a consistent modulation of perilipin 1 levels (PLIN1) suggesting that the tissue is in an equilibrium state (**Fig. 3H-I**) ^41^. The apparent contrast between the levels of PKA phospho-substrates and phospho-HSL Ser_563_, led us to hypothesize that the analysis of the signaling cascade *in vivo* could be altered by confounding factors due to tissue heterogeneity. To overcome this limitation, we take advantage of isolated mPBAs under stimulation with isoproterenol (ISO 10 μM, 16h), a well-known agonist of beta-adrenoreceptor ^42^. Western blot analysis results show that SOD1-G93A mPBAs have a defective induction of the lipolytic cascade downstream to PKA, since SOD1-G93A cells are unable to induce PKA-substrates phosphorylation under ISO treatment, in respect to wild type cell. In contrast, phosphorylation of HSL Ser_563_ appears unchanged between wild type and SOD1-G93A mPBAs under ISO stimulation (Fig. 3 J-K). This discrepancy could indicate that activation of PKA and HSL phosphorylation could be regulated by different kinetics. In fact, terminal lipolysis, assessed by the levels of PLIN1, clearly show that both cell types are competent in degrading lipid droplets under stimulation, although at the basal level, SOD1-G93A cells have a reduced lipid content and lower PLIN1 levels (**Fig. 3 J-K**), as also showed in Fig. 2 A-D. Interestingly, SOD1-G93A mPBAs clearly show an increase in SERCA1 protein expression under ISO stimulation (Fig. 3 J-K), still confirming that these cells are more prone to activate the alternative thermogenic program as suggested by the multi-omics analysis. Overall, the data shown above, clearly highlight defects in primary brown adipocytes from SOD1-G93A mice at different levels, suggesting that alterations observed with the multi-omics characterization are imputable to cell-autonomous impairments due to the diseased context.

**Figure 3:**
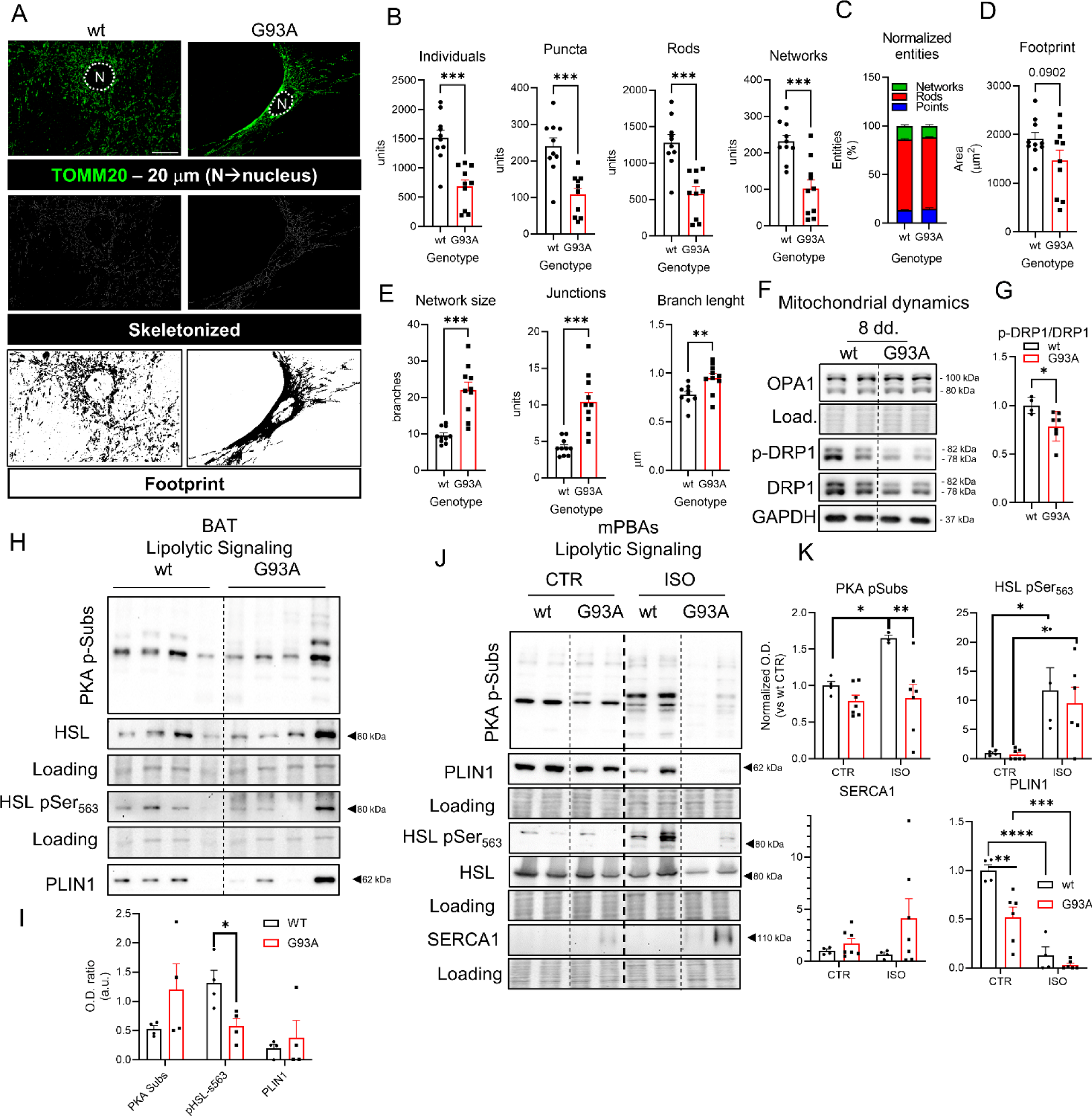
characterization of murine primary brown adipocytes from G93A mouse model. A) representative fluorescent micrographs of primary brown pre-adipocytes labelled with anti-TOMM20 and Alexafluor-488 secondary antibody. The nuclear space (N) is delimited by dotted line. Scale bar 20 μm. The skeletonized image and the footprint image were retrieved from the output of the MiNA Image J plugin. B) bar plot representing the different morphological parameters of mitochondrial individuals (individuals, rods, puncta and networks). Data are reported as mean ± SEM. Statistical significance was assessed through Student’s t-test. *** p<0.001. C) stacked bar plot representing the percentage of mitochondrial entities (puncta, rods and networks) over total. D) bar plot representing the footprint of the total mitochondrial area expressed in μm^2^. Statistical analysis was assessed through Student’s t-test. Numeric p-value was reported. E) bar plot reporting the data of the network parameters (network size, number of junctions and branch length). Data are represented as mean ± SEM. Statistical analysis was performed through Student’s t-test. ** p<0.01 *** p<0.001. F) representative western blot of proteins involved in the mitochondrial dynamics DRP-1 p-Ser_616_, DRP-1 and OPA-1. Loading control is shown as Ponceau Red staining and GAPDH protein levels. G) bar plot reporting the densitometric quantitation of DRP-1 pSer_616_ normalized versus total protein levels of DRP-1. Data are reported as mean ± SEM. Statistical analysis was performed through Student’s t-test. * p<0.05. H) representative western blot of BAT tissue lysates from wild type and SOD1-G93A mouse model. Lipolytic proteins and targets were analyzed (PKA p-Substrates, HSL, HSL p-Ser_563_, PLIN1). Ponceau Red was used as loading control. I) bar plot of densitometric analysis of western blot in panel N. Data are represented as mean ± SEM. Statistical significance was assessed by Student’s t-test. * p<0.05. J) representative western blot of mPBAs from wild type and SOD1-G93A mouse models, treated or not with isoproterenol (10 μM, 16h). Lipolytic signaling proteins (PKA p-Substrates, PLIN1, HSL p-Ser_563_ and HSL) and alternative thermogenesis (SERCA1) were analyzed. Ponceau Red staining was used as loading control. K) bar plot representing the densitometric quantitation of western blots shown in panel P. Data are presented are mean ± SEM and normalized versus wild type CTR group. Statistical significance was assessed by Two-Way ANOVA with multiple comparisons correction. * p<0.05; ** p<0.01; *** p<0.001.

### 3.4. Extracellular vesicles from SOD1-G93A BAT show alterations in size distribution and cargo

We asked whether also in ALS BAT-EVs production could be altered. To address this question, we performed EVs isolation through differential ultracentrifugation and analyzed the size distribution with Nanoparticle Tracking Analysis. Results show a positive shift in size distribution frequency and area under curve (AUC) analysis, reporting an increased EVs secretion from SOD1-G93A BAT accounting to 4.87 x 10^7^ particles/mg tissue versus 3.53 x 10^7^ particles/mg tissue of wild type BAT (**Fig. 4A-B**). To assess whether particular size range were differentially represented in SOD1-G93A BAT-EVs, we performed data normalization versus the wild type counterpart. Data reported in Figure 4C show a significant decrease of vesicles with diameter between 66-69 nm, while increasing with diameter between 213-225 nm, 325-335 nm, 355-358 nm and 462-484 nm (**Fig. 4C**). Further data analysis confirmed that wild type EVs have lower number of particles with diameter greater that 160 nm, in respect to particles ranging 0-160 nm, while SOD1-G93A particles have lower amount of particles in the 0-160 nm range and increased abundance in the > 160 nm interval (**Fig. 4D**). Overall, EVs mean diameter increases from 163.2 ± 3.080 nm in wild type to 181 ± 16.27 nm in SOD1-G93A, while modal diameter increases from 113.9 ± 1.857 nm in wild type to 126.7 ± 4.8 in SOD1-G93A EVs nm (**Fig. 4E-F**). In parallel, D10, D50 and D90 parameters respectively represent the percentage of vesicles with diameter lower than the indicated. The increase in all the three parameters strengthen the information that particles from SOD1-G93A BAT mice have an increase in size (**Fig. 4G**). To better understand whether the pathological context could also impact on the EVs cargo, we performed proteomics analysis on total EVs lysates. Although correlation plot and PCA analysis did not show a significant segregation of wild type versus SOD1-G93A samples (**Supp. Fig. 3A-B**), statistical analysis performed with Student’s t-test showed significant increase of different proteins (**Fig. 4H** and **Supp. Fig. 3C**), related to the metabolic processes (**Supp. Fig. 3D**), with particular reference to mitochondrial metabolic enzymes (**Fig. 4I**). Enrichment map analysis showed that upregulated proteins rely to TCA cycle, propanoate metabolic pathway, metabolic processes of dicarboxylic and monocarboxylic acids and regulation of ketone body metabolism (**Fig. 4J** and **Supp. Fig. 3E**). PPI network shows that all entities are interconnected, suggesting high correlation of functions of proteins secreted through BAT-EVs (**Supp. Fig. 3F**). Moreover, MCODE clustering of the PPI network recognized two main subnetworks. The MCODE1 subcluster contains all the subunits of the mitochondrial pyruvate dehydrogenase complex, namely the pyruvate dehydrogenase holoenzyme (PDHX), the dihydrolipoamide dehydrogenase (DLD) and the dihydrolipoyl transacetylase (DLAT). Moreover, also pyruvate dehydrogenase kinase (PDK1) and isocitrate dehydrogenase isoform 3A (IDH3A) are overrepresented in this cluster. MCODE subcluster 2 contains enzymes involved in fatty acid metabolism. In particular, enoyl-coenzyme A hydratase (ECH1) and hydroxyacyl-coenzyme A dehydrogenase (HADH) are involved in fatty acid beta oxidation catabolism, while propionyl-coenzyme A carboxylase isoenzyme A and B (PCCA and PCCB) are involved in fatty acids biosynthesis (**Fig. 4K**). To get further information about the differential regulation of the proteomics profile between the intracellular and the extracellular environment, we performed correlation analysis and 2D enrichment on BAT proteomics versus BAT-EVs proteomics (**Fig. 4L-M**). 2D enrichment analysis showed negative correlation between BAT and BAT-EVs proteomes, with mitochondrial proteins occupying the lower-right quadrant, which represents entities underrepresented in the intracellular milieu while being overrepresented in the EVs compartment (**Fig. 4M**). Heatmap representation of all mitochondrial proteins identified in EVs, clearly show enrichment in SOD1-G93A EVs (**Fig. 4N**). To confirm the proteomics data, we performed western blot analysis of total EVs lysates. As suggested, PDHB is one of the mostly upregulated proteins (**Fig. 4O-P**). Interestingly, also ETC complex III subunit UQCRC2 significantly increases suggesting that ejection of mitochondrial proteins through the EVs compartment could also involve proteins from the inner mitochondrial membrane (**Fig. 4O-P**). Overall, the reported data suggest a modulation of the vesicular secretory function of BAT in the SOD1-G93A context, with particular relevance to the selective secretion of mitochondrial particles.

**Figure 4:**
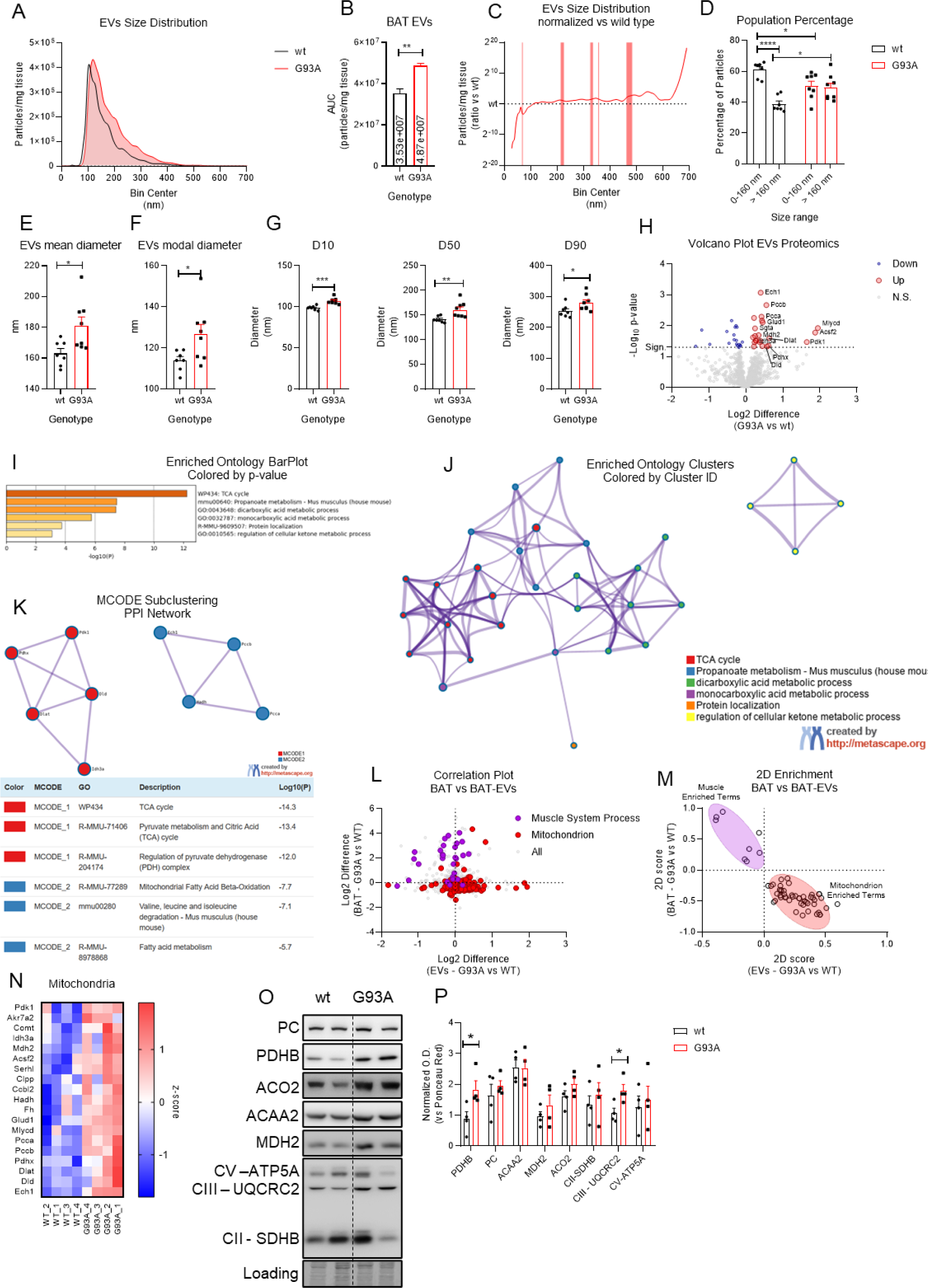
Characterization of BAT-EVs. A) distribution plot of BAT-EVs size from wt and SOD1-G93A mouse models. Data are presented as particles, normalized per unit of weight of BAT tissue. B) bar plot representing the BAT-EVs concentration calculated as area under curve (AUC) from panel A. Data are presented as mean ± SEM. Statistical significance was assessed by Student’s t-test corrected for AUC analysis. ** p<0.01. C) distribution plot of BAT-EVs size, normalized versus the wild type group. Statistical significance between size bins was estimated by multiple t-test assuming gaussian distribution and homoscedasticity, without multiple comparisons correction. Statistically significant bin ranges are highlighted in red. D) bar plot representing the quantitation of population percentage of BAT-EVs from wild type and SOD1-G93A in the 0-160 nm and >160 nm ranges. Statistical significance was assessed by Two-Way ANOVA with multiple comparisons correction. * p<0.05; *** p<0.001. E-G) bar plot representing the EVs mean and modal diameter and the D10, D50 and D90 parameters. Data are presented as mean ± SEM. Data were analyzed by Student’s t-test. * p<0.05; ** p>0.01; *** p<0.001. H) volcano plot representing the proteomics profiling of BAT-EVs from SOD1-G93A versus wild type samples. I) bar plot from Metascape analysis representing the - Log10 p value of the enrichment scores from proteomics analyses of BAT tissue and BAT-EVs integration. J) enrichment map of the integrated proteomics data of BAT tissue and BAT-EVs from wild type and SOD1-G93A mouse models. K) PPI network of MCODE clusters identified in the integrated PPI networks of proteomics data from wild type and SOD1-Gi3A mouse model BAT tissue and BAT EVs. For each cluster, the top 3 enriched terms are reported by -Log10 p-value. L) dispersion plot representing the correlation between expression values of proteins from the proteomics dataset of BAT tissue and BAT-EVs. M) dispersion plot representing the 2D-Enrichment analysis of BAT tissue and BAT-EVs. The two groups of muscle-related and mitochondrial-related entities are highlighted in purple and red, respectively. N) heatmap representing the z-scored values of mitochondrial proteins from the proteomics analysis of BAT-EVs from wild type and SOD1-G93A mouse models. O) representative western blot analysis of BAT-EVs from wild type and SOD1-G93A mouse models. mitochondrial proteins and ETC complexes were analyzed. Ponceau Red staining was used as loading control. P) bar plot representing the densitometric quantitation of western blot in panel O. data are reported as mean ± SEM of normalized OD value versus loading control. Data were analyzed with Student’s t-test. * p<0.05.

**Supplementary Figure 3:**
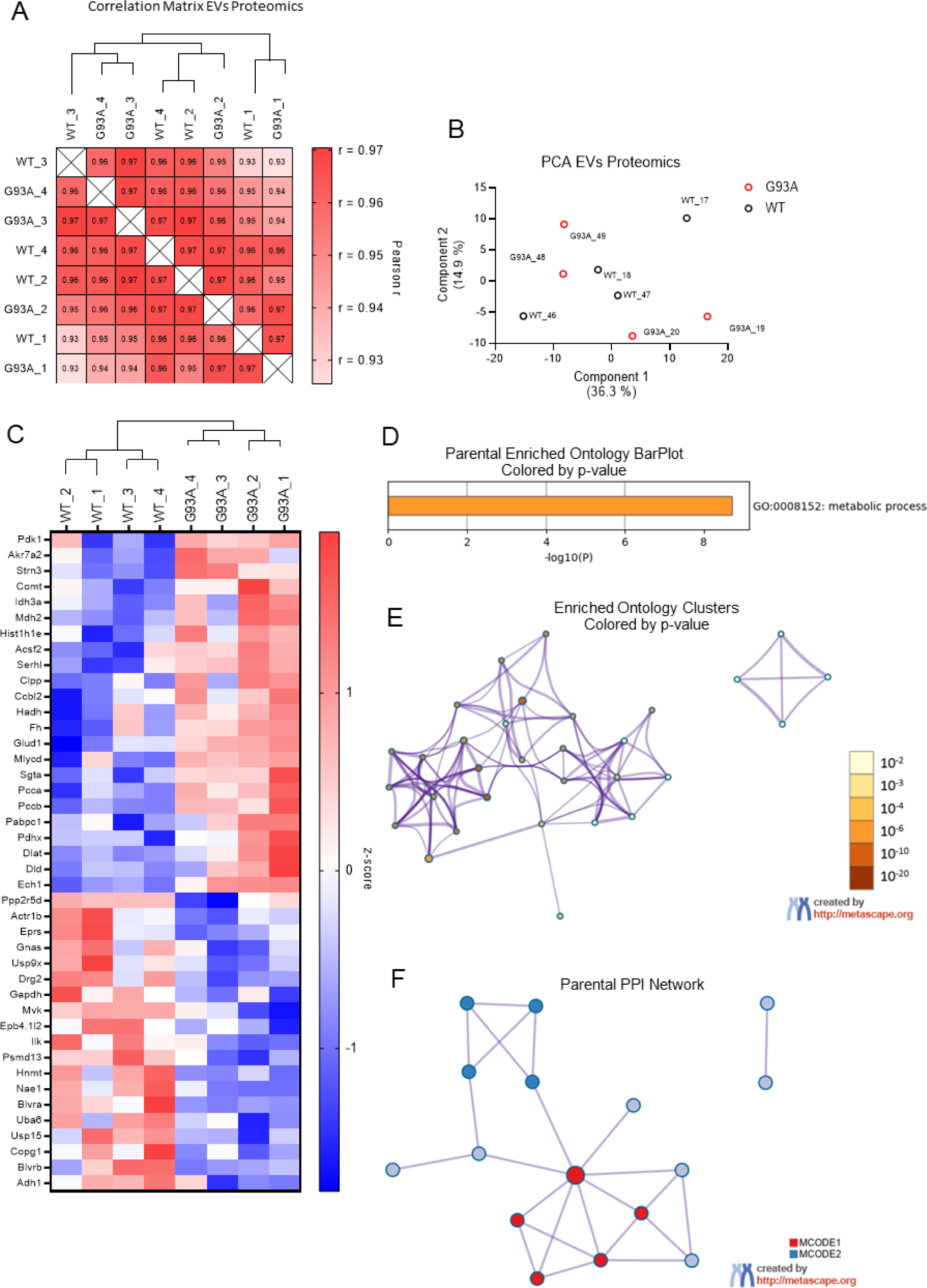
proteomics and enrichment analysis of BAT EVs supplementary information. A) heatmap representing the Pearson correlation r values for dataset in proteomics data of BAT-EVs. B) dispersion plot representing the principal component analysis of BAT EVs from wild type and G93A mouse models. C) heatmap representing the significantly regulated proteins in the proteomic analysis of BAT-EVs from wild type and G93A mouse models. Values are normalized in z-scoring and plotted as red (for up-regulated) and blue (for down-regulated). D) bar plot representing the p-value of the parental ontology category relative to the enrichment analysis in Fig. 4I. E) Enrichment map colored by p-value, relatively to the Fig. 4J. F) parental enrichment map relative to the MCODE clustering on Fig. 4K.

## 4. BAT-EVs from SOD1-G93A mice modulate differentiation and homeostasis of C2C12 murine myoblasts

With the aim of investigating the possible crosstalk between the BAT and the skeletal muscle in the ALS context, we started our investigation with a study about the capability of BAT-EVs on modulating C2C12 murine myoblasts viability. BAT-EVs from both wild type and SOD1-G93A mouse models are able to decrease C2C12 viability at a concentration over 4 μg/ml (**Fig. 4A**). Non-linear regression analysis allowed the calculation of the IC50 for both BAT-EVs types, accounting for 4.4 μg/ml for wild type and 2.6 μg/ml for SOD1-G93A BAT-EVs (**Fig. 4B**). We decided to follow up our experiments in condition of a mean dose of BAT-EVs, relatively to the IC50 calculated in **Fig. 4B** (3.5 μg/ml). We next tested the effect of BAT-EVs on C2C12 myoblasts differentiation, by treatment at T0 in differentiation medium and by analyzing the expression of myogenic transcription factors (MyoD and myogenin at 24 and 48 hours after treatment) and end-point differentiation, at 72 hours (**Fig. 4C**). Results shown in **Fig. 4 D-E** show that MyoD does not change consistently between samples, while myogenin significantly increases in samples treated with BAT-EVs from wild type animals, and such increase is not recapitulated by SOD1-G93A BAT-EVs (Fig. 4 E) suggesting that wild type BAT EVs could stimulate the early phase of myogenic differentiation. BAT-EVs from SOD1-G93A mouse models were able to inhibit C2C12 terminal differentiation, as calculated by the fusion index analysis, in respect to the samples treated with BAT-EVs from wild type mice (Fig. 4 F-G). Given that the myogenic differentiation correlates with an increase of the mitochondrial performance we performed the electron flow assay by using the Seahorse technology that assess the ETC complex enzymatic activity (Fig. 4 H). Complex I, II, and III-dependent oxygen consumption rate is increased in the wild type BAT-EVs-treated group both in respect to the NT and the SOD1-G93A BAT-EVs-treated group (Fig. 4 I). These data suggest that BAT-EVs are capable of modulating the homeostasis of murine myoblasts during the differentiation process, by stimulating the early transcriptional myogenic cascade and the mitochondrial performance. Finally, we tested the effect of BAT-EVs on differentiated C2C12-derived myotubes (Fig. 4 J). To this aim we treated myotubes with 3.5 μg/ml of BAT-EVs. Results show that SOD1-G93A BAT-EVs are capable of inducing myotube atrophy (Fig. 4 K), as assessed by the calculation of myotube diameter (Fig. 4 L). Consistently, such phenotypical alteration parallels with increased expression of E3-ubiquitin ligase Murf1 but not of Atrogin (Fig. 4 M), suggesting a different modulation of the two genes by SOD1-G93A BAT-EVs.

**Figure 4:**
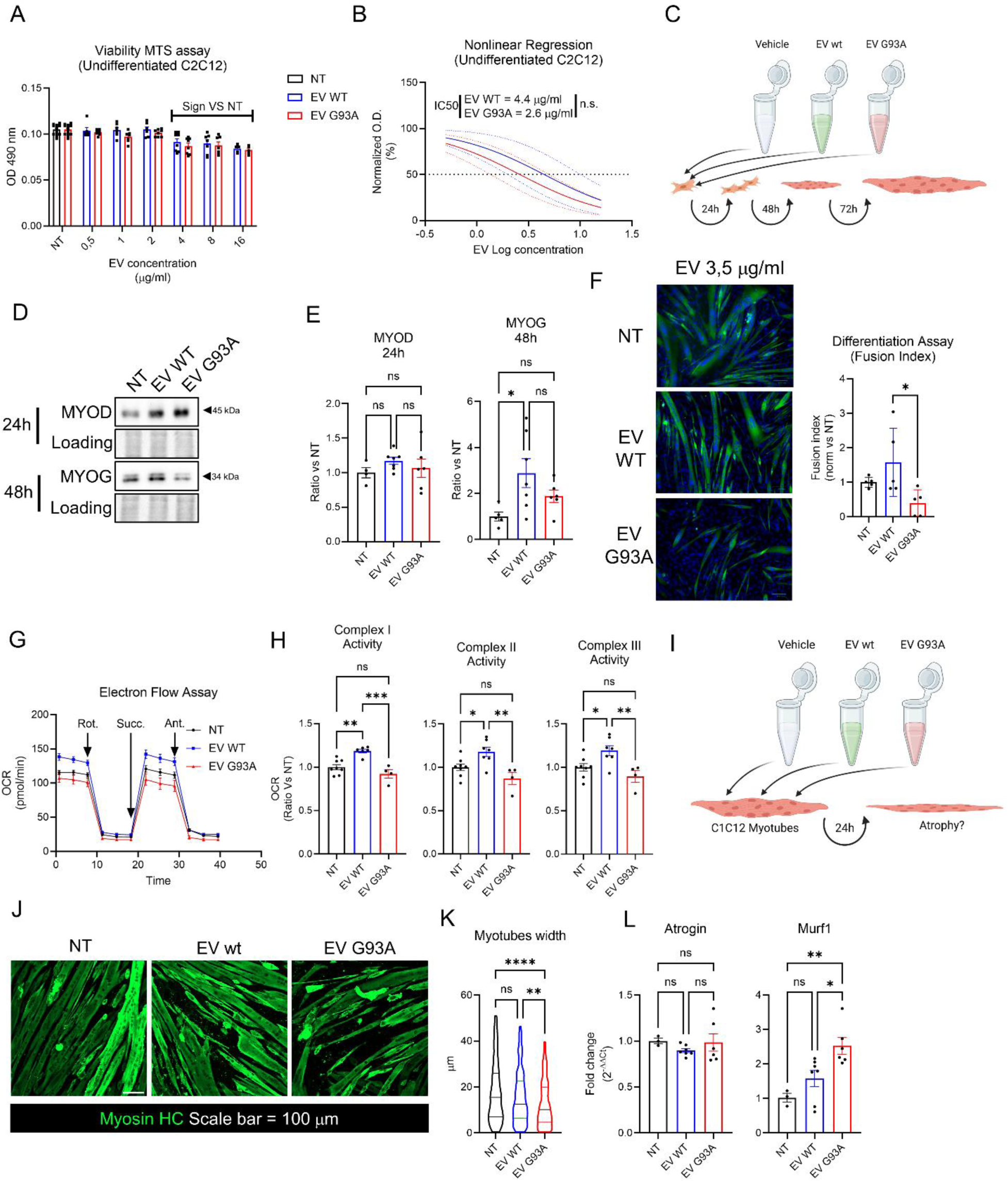
effect of BAT-EVs on C2C12 murine myoblasts. A) Bar plot representing the MTS viability assay on C2C12 myoblasts treated with BAT-EVs from wild type and SOD1-G93A mice. O.D. at 490 nm were normalized versus non-treated (NT) samples. Data are reported as mean ± SEM. Statistical significance was assessed through Two-Way ANOVA with multiple comparison correction. n = 6. B) curve fitting of data in panel A. O.D. was normalized between 0 (minimum value) and 100% (maximum value), EV concentration was logarithmized and fitting was performed through non-linear regression analysis using a dose-response inhibitory model with variable slope. Dotted lines correspond to 95% confidence intervals. IC50 was returned by the software and statistical significance was assessed by Student’s t-test. C) cartoon representing the experimental model of C2C12 murine myoblasts treated with BAT-EVs at time zero of differentiation. Differentiation process was monitored at 24, 48 and 72 hours after treatment. Artwork was made with Biorender https://www.biorender.com/. D) representative western blot of C2C12 cells at 24 and 48 hours after treatment with BAT-EVs. MyoD was assessed at 24 hours. Myogenin was assessed at 48 hours. Ponceau Red staining was used as loading control. E) bar plot representing the densitometric quantitation of western blot in panel D. Data are reported as mean ± SEM and OD of bands were normalized versus the NT sample. Statistical significance was assessed by One-Way ANOVA with multiple comparisons correction. * p < 0.05. MyoD: n_NT_ = 4; n_WT_ = 7 n_G93A_ = 6. Myogenin: n_NT_ = 5; n_WT_ = 7; n_G93A_ = 6. F) representative fluorescent micrographs of C2C12-derived myotubes at 72 hours after treatment. Sarcomeric myosin was labelled with anti-MHC (Mf-20 clone). Nuclei were counterstained with Hoechst 33342. Sclae bar 100 μm. G) bar plot representing the fusion index calculated for experiment in panel F. fusion index is expressed as percentage of myonuclei over total number of nuclei per field. Data were normalized versus NT sample. Data are reported as mean ± SEM. Statistical significance was assessed by One-Way ANOVA with multiple comparison correlations. * p< 0-05. n_NT_ = 5; n_WT_ = 5; n_G93A_ = 5. H) graph representing the oxygen consumption rate (OCR) of undifferentiated C2C12 cells treated with BAT-EVs, during the electron flow experiment in Seahorse XF assay. Mitochondria were stimulated with rotenone (complex I inhibitor), succinate (Complex II activator) and antimycin A (complex III inhibitor). Data are expressed in pmol O_2_/min. I) bar plots representing the electron transport chain complexed (ETC) activity assessed by the electro flow in panel H. data are expressed as OCR and normalized versus the NT sample. Statistical significance was assessed by the One-Way ANOVA test with multiple comparisons correction. * p<0.05 ** p<0.01 *** p< 0.001. J) cartoon representing the experimental model of C2C12-derived myotubes treated with BAT-EVs. Artwork was made with Biorender https://www.biorender.com/. K) representative fluorescent micrographs of C2C12.derived myotubes treated with BAT-EVs. Sarcomeric myosin was labelled with anti-MHC (Mf-20 clone). Scale bar = 100 μm. L) violin-plot representing the myotubes width calculated from experiment in panel K. myotubes width was calculated with an Image J dedicated plugin and expressed in μm. data derive from single myotubes analyzed from 6 different images (one per sample in the center of the well). Statistical significance was assessed through Kruskal-Wallis test for non-normal distributions with multiple comparison correction. dashed line represents median values. Dotted lines represent 1^st^ and 3^rd^ quartiles. ** p<0.01 **** p<0.0001. n_NT_ = 274; n_WT_ = 368; n_G93A_ = 318. M) bar plot representing the RT-qPCR analysis of Atrogin and Murf1 genes upon treatment with BAT-EVs for 24 hours. Data were analyzed with the 2^-ΔΔCt^ method and reported as mean ± SEM versus NT sample. Statistical significance was assessed through the One-Way ANOVA test with multiple comparison correction. * p<0.05 ** p<0.01. n_NT_ = 3; n_WT_ = 7; n_G93A_ = 6.

## 5. Discussion

Among the most peculiar clinical manifestations of ALS, thermoregulation has been one of the most debated by the scientific community. Indeed, patients can differently report about feeling hot or, conversely, being unable to warm up. In animal models, it has been shown that SOD1-G93A mice have thermoregulatory defects, being unable to maintain the body temperature upon fasting, cold exposure or morphine injection^43–45^. Considering this evidence, mouse models suggest that thermoregulation stands for a defective physiological system in ALS pathology. Consistently, thermoregulation strictly relies on the systemic energetic balance and homeostasis, shown to be altered in ALS patients and mouse models^46^. In our study, we applied a multi-omic approach for investigating the perturbations occurring into BAT from ALS SOD1-G93A mouse model at symptomatic stage. Interestingly, alternative thermogenic mechanisms are highly upregulated, and rely on the increased number of genes and proteins classically known to play an active role in the skeletal muscle homeostasis, namely, the acto-myosin complex and the SERCA pumps. Interestingly, brown adipocytes have an ontogenic link with skeletal muscle cells, since multipotent Pax3^+^/Pax7^+^/Myf5^+^ stem cells provide progenitors for both the BAT and the skeletal muscle development during embryogenesis^47^. Recently, alternative thermogenic programs were described in several reports. Aquilano and colleagues^48^ showed that dietary modulation consisting in low-protein/high-carb diet was able to induce thermogenic pathways relying on futile cycles as the upregulation of acto-myosin proteins and sarco-endoplasmic reticulum calcium ATPases (SERCA) pumps in beige adipocytes. Despite, acto-myosin complexes do not provide contractile properties to adipocytes but increase heat generation through futile cycle of ATP hydrolysis^49^. SERCA pumps are essential to control calcium homeostasis and balance between cytosol and endoplasmic reticulum reservoir. SERCAs pump back calcium into the ER lumen trough active transport, consuming ATP. Moreover, several mechanisms, as sarcolipin and phospholamban overexpression, uncouple ATP hydrolysis from calcium uptake, thus providing another futile cycle for heat dissipation^50^. These alternative thermogenic mechanisms could be important in physio-pathological situations where mitochondrial functionality is impaired, and BAT needs to redirect heat dissipation to non-mitochondrial systems. Generally, we refer to these mechanisms as “futile cycles”, that are defined as all those mechanisms that consume ATP without conferring a proper function to the cell. Adipose tissue, both white and brown, can trigger futile cycles in different conditions, such as: 1) the endogenous mitochondrial UCP1-independent uncoupling, mediated by the AAC protein in BAT ^51,52^; 2) the creatine-dependent ADP/ATP cycling mediated by both AAC protein and creatine kinase (CK) in beige fat ^53,54^; 3) the glycerolipid-free fatty acid cycle ^55,56^; 4) the glyceroneogenesis-lipid cycle, one of the very first discovered in the late 70s ^57^.

The activation of such alternative genes could be dependent on the unaltered response by UCP1 gene/proteins, revealed in our study. This is partly in contrast with some report, although this information has always been disputed. Dupuis and colleagues^43^ did not find consistent changes in UCP1 levels in BAT from SOD1-G86R mice at symptomatic stage, while, more recently, Bayer and colleagues^58^ and Ciccarone and colleagues^59^ found upregulation of UCP1 and mitochondrial proteins both in BAT and WAT from SOD1-G93A and FUS mouse models. In contrast, FUS-related alterations were not observed by Scekic-Zahrinovic and colleagues^60^ and Bayer and colleagues^58^. In 2020, Steyn and colleagues^61^, showed that BAT mass nor glucose uptake were changed in SOD1-G93A mice, from 50 to 150 days of age. Overall, all these discrepancies could likely depend on the heterogeneity of the mouse models, and stages of the disease used in the different studies. Moreover, our data show that the activation of alternative thermogenic program came with a strong downregulation of genes and proteins related to the mitochondrial compartment. We observed drastic reduction in the protein levels of pyruvate dehydrogenase isoform A and B, enzymes involved in the critical step of conversion of pyruvate to acetyl-CoA in the mitochondria, for entering the TCA cycle. PDH activity is important for maintaining energy homeostasis in ALS, as knockdown of pyruvate dehydrogenase kinase 2 (PDK2), that inhibits PDH, is able to support mitochondrial metabolism in SOD1-G93A rat model, by ameliorating the interplay between astrocytes and motor neurons ^62^. In light of this, PDH expression could represent a hub for the regulation of energy metabolism in ALS, also in peripheral tissues.

In the present study, we showed that *ex vivo* cultured mPBAs have differentiation and metabolic defects, strengthening the idea that the ALS-related feature is cell-autonomous compared to the tissue microenvironment. The decrease in adipocyte area is partly in accordance with Ciccarone et al.^59^, that showed reduction in adipocyte area in both BAT and subcutaneous WAT at the end-stage (150 d.p.p.). In contrast, they showed upregulation on UCP1 protein and mitochondrial markers. This discrepancy could likely depend on the different stage of the disease. The modulation of terminal adipogenesis correlates with the differential modulation of PPARγ isoforms. Indeed, in our study, mPBAs from SOD1-G93A mouse model have impaired expression of the PPARγ2 isoform (PPARγ2). PPARγ2 is highly expressed in browning-resistant fat depots, strengthening the information that brown adipocytes from SOD1-G93A mice has an impairment in the induction of the canonical thermogenic program^63^. The differentiation defects came with metabolic defects in mPBAs from SOD1-G93A mouse model. Metabolic defects are a typical hallmark of ALS in different tissues and cell types. In particular, the skeletal muscle of ALS mouse model shows alteration in fiber types (glycolytic-to-oxidative switch) and in mitochondrial functionalities, that predate the onset of motor symptoms ^64–67^. Notably, mitochondrial metabolism alterations have also been observed in ALS patients’ skeletal muscle samples ^67,68^. Also, the neuronal compartment has metabolic defects in the ALS context. In mouse model, both spinal and cortical neurons have hampered basal respiration, maximal respiration, spare capacity and ATP production ^69^. This alteration has been linked to defects in the autophagic process and in signaling pathways controlling energy homeostasis ^69,70^. Adipose tissue has been poorly investigated in ALS; different reports highlight that also white adipose tissue have metabolic defects. Steyn et al, showed that epididymal AT from SOD1-G93A mice have increased lipolytic efflux and reduced fat mass, also correlated with reduced plasmatic leptin concentration ^61^. According to the lipolytic process, in the present work we showed that BAT have lower activation of the lipolytic signaling under control of PKA, while mPBAs are unable to properly respond to the beta-adrenergic stimulation. Importantly, metabolic homeostasis can be rewired both in vitro and in vivo, correlating with amelioration of tissue structure and ultrastructure as well as animal motor symptoms ^64,69,71^. Moreover, in our study we presented a characterization of the mitochondrial dynamics in primary pre-adipocytes extracted from BAT of wild type and SOD1-G93A mice. Our analyses highlighted that SOD1-G93A pre-adipocytes have a more fuse mitochondrial network, with a decreased number of individuals and increased network size. This is in favor of the role of mitochondrial dynamics in supporting the metabolism and functionality of BAT. Indeed, human brown adipocytes typically exhibit fragmented mitochondrial morphology, which supports higher levels of catabolic processes, uncoupled respiration, and thermogenesis^72^. The interplay between mitochondrial fission and increased UCP1 expression may enhance thermogenesis in these cells. Several key proteins regulate the processes of mitochondrial fission and fusion. Dynamin-related protein-1 (DRP1), a cytoplasmic GTPase, plays a crucial role in this regulation; inhibition of DRP1 disrupts adrenaline-induced thermogenesis, underscoring its importance^72^. Exposure to cold triggers norepinephrine secretion, which leads to rapid mitochondrial fragmentation via PKA-dependent phosphorylation of DRP1, accompanied by lipolysis and UCP1 induction^73^. In beige adipocytes derived from human mesenchymal stem cells, DRP1-mediated fission enhances uncoupling activity^72^. Additionally, brown fat-specific deletion of the inner mitochondrial membrane fusion protein optic atrophy 1 (OPA1) results in mitochondrial dysfunction and impaired cold-induced thermogenesis^74^. Recent findings suggest that OPA1-dependent fumarate accumulation promotes cell-autonomous adipocyte browning^75^. Given this body of evidence, we speculate here that the mitochondrial defects highlighted by the functional analysis of extracellular flex and lipolysis competence, could be strictly linked to the abnormal mitochondrial dynamics here observed. In our mind, the excessive fusion of the mitochondrial network is dependent on the decreased phosphorylation of DRP-1, while OPA-1 protein remains unchanged. Possibly, alteration of signaling network (e.g. PKA-dependent) could lead to decreased mitochondrial fragmentation and consequent lower metabolic competence.

Extracellular vesicles (EV) have been shown to play a prominent role in ALS since they could be vehicle of misfolded proteins such as TDP-43, FUS, and SOD1 either and could represent an optimal biomarker for ALS diagnosis and/or prognosis, to monitor the disease progression or to develop new therapeutical approaches ^76–78^. The importance of studying EVs in ALS pathology is based on the fact that these vesicles are able to cross the blood brain barrier ^79^ it is therefore not to be excluded that also EVs from peripheral organs pivotal involved in the ALS phenotype. In the case of EVs from peripheral organs, adipose-derived stem cells (ADSCs) and mesenchymal stem cells (MSCs) have been investigated ^80–82^. Interestingly, AT-derived EVs exert a neuroprotective and anti-neuroinflammatory effect both in *in vitro* and *in vivo* models, not only in ALS, but also in other neurodegenerative disease ^83–87^. In particular, regarding ALS, AT-derived EVs are able to normalize the levels of aggregated SOD1 protein and reactivate CREB and PGC1alpha pathways, that ameliorate the mitochondrial phenotype ^88^. Moreover, AT-derived EVs can also be repeatedly administered to the SOD1-G93A mouse model, both intravenously and intranasally, being safe for the animal and able to improve motor performance and protecting both nerves and muscles from degeneration ^86^. In WAT and heart tissue mitochondrial extrusion and the interplay with immune system cells was pivotal for maintaining metabolic and inflammatory homeostasis^89–91^. Recently, our group has taken part of a study showing that physiological stimuli, as cold stress, induce mitochondrial ROS damage in BAT, which activates an alternative recycling pathway of damaged mitochondria through extracellular vesicle release. Moreover, EVs from BAT of cold-exposed mice, had oxidized and carbonylated proteins, proxy of intracellular ROS damage. Interestingly, such EV population was capable of altering the metabolism of recipient cells, and their removal from tissue-resident macrophages was necessary for the homeostatic regulation of thermogenesis^20^. Here we showed that BAT-EVs have different effect on the homeostasis of an in vitro model of myogenic cells, namely the murine C2C12 myoblasts. The interplay between BAT and skeletal muscle has been already documented, although preliminary and contradictory, in part. The evidence in support of a positive role of AT-derived EVs, show that adipocytes subjected to mitochondrial stress release EVs containing respiratory competent mitochondrial particles that can be taken-up by cardiomyocytes to support cardiac cells in the ischemic/reperfusion injury model ^92^. On the contrary, AT-derived EVs could also be a source of cytokines that can regulate skeletal muscle homeostasis at different levels, perturbing satellite cells activation, myoblasts proliferation/differentiation and protein catabolism as well. Such effects could be positive or detrimental on the skeletal muscle cell, depending on the type of cytokine and on the systemic status of the individual (lean/obese, sedentary/exercised, young/aged) ^93^. Among these effectors, Resistin has been recently found to be released by AT also through EVs ^94^ and was shown to alter the fusion process of differentiating myoblasts via the activation of the NF-kB pathway or by reducing the expression of desmin and myogenin ^95–97^. Another relevant cargo of EVs are miRNA. Although we did not consider to examine mRNAs alteration in the present work, it has been documented that adipocytes can also release miRNAs (miR-27a, miR-130b), via EVs, that can produce insulin resistance and regulated lipid metabolism through PPARγ and PGC1α inhibition ^98,99^. An interesting point about the effect of BAT-EVs in our experimental system is the detrimental effect of SOD1-G93A EVs on the mitochondrial functionality, with particular relevance to the alteration in Complex I (NDUFB8) activity. Our group have already shown that the skeletal muscle from SOD1-G93A mouse model have impairment of Complex I activity ^64^. In particular, results showed an early impairment of Complex I activity in the skeletal muscle of SOD1-G93A mice. This impairment in Complex I activity has also been noted in muscle biopsies from patients with sporadic ALS^100^. Functional alterations in Complex I were detected in gastrocnemius muscle isolated from SOD1-G93A mice at 7 weeks of age^101^. These findings collectively support the hypothesis that Complex I activity is a selective target of various ALS-related proteins in both ALS patients^102^ and preclinical models^103,104^. In the present work, we showed that Complex I protein induction if promoted at thew earliest phases of myogenic differentiation by wild type EVs, while the same induction is not provided by SOD1-G93A EVs. In parallel, SOD1-G93A EVs dampen the enzymatic activity, as shown by the electron flow assay. The induction of mitochondrial biogenesis is a key feature of myogenic differentiation ^105,106^ This is in accordance with the role known for Complex I in promoting myogenesis. It has been recently shown that Complex I is able to promote myoblasts differentiation through a mechanism involving p53 acetylation and the regulation of the NAD+/NADH ratio ^107^. Inhibition of Complex I activity also lead to alteration of the cell cycle progression and alteration of the terminal differentiation^108^. Here we speculate that Complex I could be a hub in the regulation of myoblasts homeostasis in ALS, and that this complex could be one of the main targets of pathologic EVs observed in our mouse model of ALS.

Consistently, we did not only show that BAT-EVs can modulate myoblast differentiation, but we also provided data that SOD1-G93A EVs can induce atrophy on C2C12-derived myotubes. It has been shown that BAT can secrete the myostatin protein upon loss of the transcription factor IRF4, and that the IRF4 expression can be modulated by interfering with BAT thermogenic activation ^109^. Interestingly, induction of myostatin expression could also be achieved by pharmacological treatment with simvastatin ^110^. In parallel, also beneficial effects of AT-derived EVs on the skeletal muscle trophism ^111^. By using an hindlimb ischemia model, it was shown that EV derived from AT-mesenchymal stem cells (injected intravenous of intramuscular) are able to protect the muscle from the ischemic degeneration and ameliorate the regeneration trough increased expression of MyoD, Myf5 and Pax7^112^ In the case of ALS, it needs to be further investigated whether also BAT-EVs could play a prominent role in regulating the disease outcome and/or development. Here, we speculate that BAT-EVs from wild type or SOD1-G93A models could have different biological properties on tissue and cells affected by the ALS phenotype, as motor neurons, skeletal muscle myofibers and/or stem cells, but also on microglial cells and astrocytes. While BAT-EVs from healthy animal could play a neuroprotective role, this could be not the case of SOD1-G93A model, as we have shown here that their proteomic profile undergoes a radical transformation.

## 6. Conclusions and limitations

Overall, our study pointed out seeding information about the alteration of the BAT in the ALS context. We showed that this tissue undergoes a drastic molecular rearrangement at the symptomatic stage of the mouse model disease, and that such alterations came with metabolic, phenotypical and secretory alterations of the brown adipocytes. The bioactive role of BAT-derived EVs on the modulation of the homeostasis of C2C12 myoblasts and myotubes are suggestive that BAT could also play an active role in skeletal muscle degeneration observe in ALS. Of course, the comparison of SOD1-G93A model with other mouse models (FUS, TDP-43) is needed, since it should be cleared whether our findings could be general of the ALS disease or are specifical for the SOD1-mutation. Further investigation could be made to understand whether BAT could present pathological feature also earlier on the symptom onset and if it could represent a putative target organ or to have a prognostic/diagnostic significance.

## 7. Author contributions

**Marco Rosina**: Conceptualization, Methodology, Validation, Formal analysis, Investigation, Resources, Data curation, Writing – Original Draft, Writing – Review & Editing, Visualization, Project Administration, Funding Acquisition. **Silvia Scaricamazza, Flaminia Riggio, Gianmarco Fenili, Flavia Giannessi, Alessandro Matteocci, Valentina Nesci, Illari Salvatori**: Methodology, Investigation, Data curation. **Daniela F. Angelini**: Supervision, Writing – Review & Editing. **Valerio Chiurchiù**: Supervision, Data curation, Writing – Review & Editing. **Katia Aquilano, Daniele Lettieri Barbato**: Conceptualization, Supervision, Writing – Review & Editing. **Nicola Biagio Mercuri**: Supervision, Project Administration. **Cristiana Valle, Alberto Ferri**: Conceptualization, Supervision, Funding Acquisition, Writing – Review & Editing.

## Acknowledgements and funding

This work has been supported by Starting Grant Ricerca Finalizzata 2019 "SG-2019-12368589 " to M.R.; PRIN 2022 "20223RXEEC", Ricerca Finalizzata 2019 "RF-2019-12369105”, “CNR IFT DBA.AD005.225-NUTRAGE-FOE 2021” to A.F.; PRIN 2022 "20222KSN2N" to C.V. Artworks were prepared with Biorender https://www.biorender.com/.

## 9. Data availability

All data and reagents will be made available upon reasonable request to the Lead Contact/Corresponding Author Dr. Marco Rosina. The mass spectrometry proteomics data have been deposited to the ProteomeXchange Consortium via the PRIDE ^113^ partner repository with the dataset identifier PXD054147. The RNA-Seq data are available at GEO identifier GSE273052.

## References

1. Kiernan, M.C., Vucic, S., Cheah, B.C., Turner, M.R., Eisen, A., Hardiman, O., Burrell, J.R., and Zoing, M.C. (2011). Amyotrophic lateral sclerosis. The Lancet 377, 942–955. 10.1016/S0140-6736(10)61156-7.

2. Swinnen, B., and Robberecht, W. (2014). The phenotypic variability of amyotrophic lateral sclerosis. Nat. Rev. Neurol. 10, 661–670. 10.1038/nrneurol.2014.184.

3. Taylor, J.P., Brown, R.H., and Cleveland, D.W. (2016). Decoding ALS: from genes to mechanism. Nature 539, 197–206. 10.1038/nature20413.

4. Lattante, S., Ciura, S., Rouleau, G.A., and Kabashi, E. (2015). Defining the genetic connection linking amyotrophic lateral sclerosis (ALS) with frontotemporal dementia (FTD). Trends Genet. 31, 263–273. 10.1016/j.tig.2015.03.005.

5. Al-Chalabi, A., van den Berg, L.H., and Veldink, J. (2017). Gene discovery in amyotrophic lateral sclerosis: implications for clinical management. Nat. Rev. Neurol. 13, 96–104. 10.1038/nrneurol.2016.182.

6. Ji, A.-L., Zhang, X., Chen, W.-W., and Huang, W.-J. (2017). Genetics insight into the amyotrophic lateral sclerosis/frontotemporal dementia spectrum. J. Med. Genet. 54, 145–154. 10.1136/jmedgenet-2016-104271.

7. Suzuki, N., Nishiyama, A., Warita, H., and Aoki, M. (2023). Genetics of amyotrophic lateral sclerosis: seeking therapeutic targets in the era of gene therapy. J. Hum. Genet. 68, 131–152. 10.1038/s10038-022-01055-8.

8. Desport, J.C., Preux, P.M., Truong, T.C., Vallat, J.M., Sautereau, D., and Couratier, P. (1999). Nutritional status is a prognostic factor for survival in ALS patients. Neurology 53, 1059–1063. 10.1212/wnl.53.5.1059.

9. Vaisman, N., Lusaus, M., Nefussy, B., Niv, E., Comaneshter, D., Hallack, R., and Drory, V.E. (2009). Do patients with amyotrophic lateral sclerosis (ALS) have increased energy needs? J. Neurol. Sci. 279, 26–29. 10.1016/j.jns.2008.12.027.

10. Desport, J.C., Preux, P.M., Truong, C.T., Courat, L., Vallat, J.M., and Couratier, P. (2000). Nutritional assessment and survival in ALS patients. Amyotroph. Lateral Scler. Mot. Neuron Disord. Off. Publ. World Fed. Neurol. Res. Group Mot. Neuron Dis. 1, 91–96. 10.1080/14660820050515386.

11. Kasarskis, E.J., Berryman, S., Vanderleest, J.G., Schneider, A.R., and McClain, C.J. (1996). Nutritional status of patients with amyotrophic lateral sclerosis: relation to the proximity of death. Am. J. Clin. Nutr. 63, 130–137. 10.1093/ajcn/63.1.130.

12. Kühnlein, P., Gdynia, H.-J., Sperfeld, A.-D., Lindner-Pfleghar, B., Ludolph, A.C., Prosiegel, M., and Riecker, A. (2008). Diagnosis and treatment of bulbar symptoms in amyotrophic lateral sclerosis. Nat. Clin. Pract. Neurol. 4, 366–374. 10.1038/ncpneuro0853.

13. Desport, J.C., Preux, P.M., Magy, L., Boirie, Y., Vallat, J.M., Beaufrère, B., and Couratier, P. (2001). Factors correlated with hypermetabolism in patients with amyotrophic lateral sclerosis. Am. J. Clin. Nutr. 74, 328–334. 10.1093/ajcn/74.3.328.

14. Desport, J.-C., Torny, F., Lacoste, M., Preux, P.-M., and Couratier, P. (2005). Hypermetabolism in ALS: correlations with clinical and paraclinical parameters. Neurodegener. Dis. 2, 202–207. 10.1159/000089626.

15. Dupuis, L., Corcia, P., Fergani, A., Gonzalez De Aguilar, J.-L., Bonnefont-Rousselot, D., Bittar, R., Seilhean, D., Hauw, J.-J., Lacomblez, L., Loeffler, J.-P., et al. (2008). Dyslipidemia is a protective factor in amyotrophic lateral sclerosis. Neurology 70, 1004– 1009. 10.1212/01.wnl.0000285080.70324.27.

16. Barneda, D., and Christian, M. (2017). Lipid droplet growth: regulation of a dynamic organelle. Curr. Opin. Cell Biol. 47, 9–15. 10.1016/j.ceb.2017.02.002.

17. Heinonen, S., Jokinen, R., Rissanen, A., and Pietiläinen, K.H. (2020). White adipose tissue mitochondrial metabolism in health and in obesity. Obes. Rev. Off. J. Int. Assoc. Study Obes. 21, e12958. 10.1111/obr.12958.

18. Cohen, P., and Kajimura, S. (2021). The cellular and functional complexity of thermogenic fat. Nat. Rev. Mol. Cell Biol. 22, 393–409. 10.1038/s41580-021-00350-0.

19. Roesler, A., and Kazak, L. (2020). UCP1-independent thermogenesis. Biochem. J. 477, 709–725. 10.1042/BCJ20190463.

20. Rosina, M., Ceci, V., Turchi, R., Chuan, L., Borcherding, N., Sciarretta, F., Sanchez-Diaz, M., Tortolici, F., Karlinsey, K., Chiurchiu, V., et al. (2022). Ejection of damaged mitochondria and their removal by macrophages ensure efficient thermogenesis in brown adipose tissue. Cell Metab. 34, 533–548.e12. 10.1016/j.cmet.2022.02.016.

21. Camino, T., Lago-Baameiro, N., Sueiro, A., Bravo, S.B., Couto, I., Santos, F.F., Baltar, J., Casanueva, F.F., and Pardo, M. (2022). Brown Adipose Tissue Sheds Extracellular Vesicles That Carry Potential Biomarkers of Metabolic and Thermogenesis Activity Which Are Affected by High Fat Diet Intervention. Int. J. Mol. Sci. 23, 10826. 10.3390/ijms231810826.

22. Xu, J., Cui, L., Wang, J., Zheng, S., Zhang, H., Ke, S., Cao, X., Shi, Y., Li, J., Zen, K., et al. (2023). Cold-activated brown fat-derived extracellular vesicle-miR-378a-3p stimulates hepatic gluconeogenesis in male mice. Nat. Commun. 14, 5480. 10.1038/s41467-023-41160-6.

23. Cadete, V.J.J., Deschênes, S., Cuillerier, A., Brisebois, F., Sugiura, A., Vincent, A., Turnbull, D., Picard, M., McBride, H.M., and Burelle, Y. (2016). Formation of mitochondrial-derived vesicles is an active and physiologically relevant mitochondrial quality control process in the cardiac system. J. Physiol. 594, 5343–5362. 10.1113/JP272703.

24. Clement, E., Lazar, I., Attané, C., Carrié, L., Dauvillier, S., Ducoux-Petit, M., Esteve, D., Menneteau, T., Moutahir, M., Le Gonidec, S., et al. (2020). Adipocyte extracellular vesicles carry enzymes and fatty acids that stimulate mitochondrial metabolism and remodeling in tumor cells. EMBO J. 39, e102525. 10.15252/embj.2019102525.

25. Xiong, Y., Soumillon, M., Wu, J., Hansen, J., Hu, B., van Hasselt, J.G.C., Jayaraman, G., Lim, R., Bouhaddou, M., Ornelas, L., et al. (2017). A Comparison of mRNA Sequencing with Random Primed and 3′-Directed Libraries. Sci. Rep. 7, 14626. 10.1038/s41598-017-14892-x.

26. BBMap (2023). SourceForge. https://sourceforge.net/projects/bbmap/.

27. Dobin, A., Davis, C.A., Schlesinger, F., Drenkow, J., Zaleski, C., Jha, S., Batut, P., Chaisson, M., and Gingeras, T.R. (2013). STAR: ultrafast universal RNA-seq aligner. Bioinforma. Oxf. Engl. 29, 15–21. 10.1093/bioinformatics/bts635.

28. Anders, S., Pyl, P.T., and Huber, W. (2015). HTSeq--a Python framework to work with high-throughput sequencing data. Bioinforma. Oxf. Engl. 31, 166–169. 10.1093/bioinformatics/btu638.

29. Love, M.I., Huber, W., and Anders, S. (2014). Moderated estimation of fold change and dispersion for RNA-seq data with DESeq2. Genome Biol. 15, 550. 10.1186/s13059-014-0550-8.

30. Zhou, Y., Zhou, B., Pache, L., Chang, M., Khodabakhshi, A.H., Tanaseichuk, O., Benner, C., and Chanda, S.K. (2019). Metascape provides a biologist-oriented resource for the analysis of systems-level datasets. Nat. Commun. 10, 1523. 10.1038/s41467-019-09234-6.

31. Ngo, J., Benador, I.Y., Brownstein, A.J., Vergnes, L., Veliova, M., Shum, M., Acín-Pérez, R., Reue, K., Shirihai, O.S., and Liesa, M. (2021). Isolation and functional analysis of peridroplet mitochondria from murine brown adipose tissue. STAR Protoc. 2, 100243. 10.1016/j.xpro.2020.100243.

32. Schindelin, J., Arganda-Carreras, I., Frise, E., Kaynig, V., Longair, M., Pietzsch, T., Preibisch, S., Rueden, C., Saalfeld, S., Schmid, B., et al. (2012). Fiji: an open-source platform for biological-image analysis. Nat. Methods 9, 676–682. 10.1038/nmeth.2019.

33. Reggio, A., Rosina, M., Palma, A., Cerquone Perpetuini, A., Petrilli, L.L., Gargioli, C., Fuoco, C., Micarelli, E., Giuliani, G., Cerretani, M., et al. (2020). Adipogenesis of skeletal muscle fibro/adipogenic progenitors is affected by the WNT5a/GSK3/beta-catenin axis. Cell Death Differ. 27, 2921–2941. 10.1038/s41418-020-0551-y.

34. Reggio, A., Rosina, M., Krahmer, N., Palma, A., Petrilli, L.L., Maiolatesi, G., Massacci, G., Salvatori, I., Valle, C., Testa, S., et al. (2020). Metabolic reprogramming of fibro/adipogenic progenitors facilitates muscle regeneration. Life Sci. Alliance 3, e202000646. 10.26508/lsa.202000660.

35. Reggio, A., Spada, F., Rosina, M., Massacci, G., Zuccotti, A., Fuoco, C., Gargioli, C., Castagnoli, L., and Cesareni, G. (2019). The immunosuppressant drug azathioprine restrains adipogenesis of muscle Fibro/Adipogenic Progenitors from dystrophic mice by affecting AKT signaling. Sci. Rep. 9, 4360. 10.1038/s41598-019-39538-y.

36. Palma, A., Cerquone Perpetuini, A., Ferrentino, F., Fuoco, C., Gargioli, C., Giuliani, G., Iannuccelli, M., Licata, L., Micarelli, E., Paoluzi, S., et al. (2019). Myo-REG: A Portal for Signaling Interactions in Muscle Regeneration. Front. Physiol. 10, 1216. 10.3389/fphys.2019.01216.

37. Carpenter, A.E., Jones, T.R., Lamprecht, M.R., Clarke, C., Kang, I.H., Friman, O., Guertin, D.A., Chang, J.H., Lindquist, R.A., Moffat, J., et al. (2006). CellProfiler: image analysis software for identifying and quantifying cell phenotypes. Genome Biol. 7, R100. 10.1186/gb-2006-7-10-r100.

38. Jurberg, A.D., Gomes, G., Seixas, M.R., Mermelstein, C., and Costa, M.L. (2023). Improving quantification of myotube width and nuclear/cytoplasmic ratio in myogenesis research. Comput. Methods Programs Biomed. 230, 107354. 10.1016/j.cmpb.2023.107354.

39. Valente, A.J., Maddalena, L.A., Robb, E.L., Moradi, F., and Stuart, J.A. (2017). A simple ImageJ macro tool for analyzing mitochondrial network morphology in mammalian cell culture. Acta Histochem. 119, 315–326. 10.1016/j.acthis.2017.03.001.

40. Cox, J., and Mann, M. (2012). 1D and 2D annotation enrichment: a statistical method integrating quantitative proteomics with complementary high-throughput data. BMC Bioinformatics 13 *Suppl 16*, S12. 10.1186/1471-2105-13-S16-S12.

41. Sztalryd, C., and Brasaemle, D.L. (2017). The perilipin family of lipid droplet proteins: Gatekeepers of intracellular lipolysis. Biochim. Biophys. Acta Mol. Cell Biol. Lipids 1862, 1221–1232. 10.1016/j.bbalip.2017.07.009.

42. Miller, C.N., Yang, J.-Y., England, E., Yin, A., Baile, C.A., and Rayalam, S. (2015). Isoproterenol Increases Uncoupling, Glycolysis, and Markers of Beiging in Mature 3T3-L1 Adipocytes. PLOS ONE 10, e0138344. 10.1371/journal.pone.0138344.

43. Dupuis, L., Oudart, H., René, F., Gonzalez De Aguilar, J.-L., and Loeffler, J.-P. (2004). Evidence for defective energy homeostasis in amyotrophic lateral sclerosis: Benefit of a high-energy diet in a transgenic mouse model. Proc. Natl. Acad. Sci. U. S. A. 101, 11159–11164. 10.1073/pnas.0402026101.

44. Dupuis, L., Petersen, Å., and Weydt, P. (2018). Thermoregulation in amyotrophic lateral sclerosis. Handb. Clin. Neurol. 157, 749–760. 10.1016/B978-0-444-64074-1.00046-X.

45. Kandinov, B., Korczyn, A.D., Rabinowitz, R., Nefussy, B., and Drory, V.E. (2011). Autonomic impairment in a transgenic mouse model of amyotrophic lateral sclerosis. Auton. Neurosci. 159, 84–89. 10.1016/j.autneu.2010.09.002.

46. Dupuis, L., Pradat, P.-F., Ludolph, A.C., and Loeffler, J.-P. (2011). Energy metabolism in amyotrophic lateral sclerosis. Lancet Neurol. 10, 75–82. 10.1016/S1474-4422(10)70224-6.

47. Jung, S.M., Sanchez-Gurmaches, J., and Guertin, D.A. (2019). Brown Adipose Tissue Development and Metabolism. Handb. Exp. Pharmacol. 251, 3–36. 10.1007/164_2018_168.

48. Aquilano, K., Sciarretta, F., Turchi, R., Li, B.-H., Rosina, M., Ceci, V., Guidobaldi, G., Arena, S., D’Ambrosio, C., Audano, M., et al. (2020). Low-protein/high-carbohydrate diet induces AMPK-dependent canonical and non-canonical thermogenesis in subcutaneous adipose tissue. Redox Biol. 36, 101633. 10.1016/j.redox.2020.101633.

49. Guarnieri, A.R., Benson, T.W., and Tranter, M. (2022). Calcium cycling as a mediator of thermogenic metabolism in adipose tissue. Mol. Pharmacol. 102, 51–59. 10.1124/molpharm.121.000465.

50. Gorski, P.A., Ceholski, D.K., and Young, H.S. (2017). Structure-Function Relationship of the SERCA Pump and Its Regulation by Phospholamban and Sarcolipin. Adv. Exp. Med. Biol. 981, 77–119. 10.1007/978-3-319-55858-5_5.

51. Bertholet, A.M., Chouchani, E.T., Kazak, L., Angelin, A., Fedorenko, A., Long, J.Z., Vidoni, S., Garrity, R., Cho, J., Terada, N., et al. (2019). H+ transport is an integral function of the mitochondrial ADP/ATP carrier. Nature 571, 515–520. 10.1038/s41586-019-1400-3.

52. Kreiter, J., Rupprecht, A., Škulj, S., Brkljača, Z., Žuna, K., Knyazev, D.G., Bardakji, S., Vazdar, M., and Pohl, E.E. (2021). ANT1 Activation and Inhibition Patterns Support the Fatty Acid Cycling Mechanism for Proton Transport. Int. J. Mol. Sci. 22, 2490. 10.3390/ijms22052490.

53. Kazak, L., Chouchani, E.T., Jedrychowski, M.P., Erickson, B.K., Shinoda, K., Cohen, P., Vetrivelan, R., Lu, G.Z., Laznik-Bogoslavski, D., Hasenfuss, S.C., et al. (2015). A creatine-driven substrate cycle enhances energy expenditure and thermogenesis in beige fat. Cell 163, 643–655. 10.1016/j.cell.2015.09.035.

54. Sun, Y., Rahbani, J.F., Jedrychowski, M.P., Riley, C.L., Vidoni, S., Bogoslavski, D., Hu, B., Dumesic, P.A., Zeng, X., Wang, A.B., et al. (2021). Mitochondrial TNAP controls thermogenesis by hydrolysis of phosphocreatine. Nature 593, 580–585. 10.1038/s41586-021-03533-z.

55. Mottillo, E.P., Balasubramanian, P., Lee, Y.-H., Weng, C., Kershaw, E.E., and Granneman, J.G. (2014). Coupling of lipolysis and de novo lipogenesis in brown, beige, and white adipose tissues during chronic β3-adrenergic receptor activation. J. Lipid Res. 55, 2276–2286. 10.1194/jlr.M050005.

56. Veliova, M., Ferreira, C.M., Benador, I.Y., Jones, A.E., Mahdaviani, K., Brownstein, A.J., Desousa, B.R., Acín-Pérez, R., Petcherski, A., Assali, E.A., et al. (2020). Blocking mitochondrial pyruvate import in brown adipocytes induces energy wasting via lipid cycling. EMBO Rep. 21, e49634. 10.15252/embr.201949634.

57. Feldman, D., and Hirst, M. (1978). Glucocorticoids and regulation of phosphoenolpyruvate carboxykinase activity in rat brown adipose tissue. Am. J. Physiol. 235, E197–202. 10.1152/ajpendo.1978.2352.E197.

58. Bayer, H., Lang, K., Buck, E., Higelin, J., Barteczko, L., Pasquarelli, N., Sprissler, J., Lucas, T., Holzmann, K., Demestre, M., et al. (2017). ALS-causing mutations differentially affect PGC-1α expression and function in the brain vs. peripheral tissues. Neurobiol. Dis. 97, 36–45. 10.1016/j.nbd.2016.11.001.

59. Ciccarone, F., Castelli, S., Lazzarino, G., Scaricamazza, S., Mangione, R., Bernardini, S., Apolloni, S., D’Ambrosi, N., Ferri, A., and Ciriolo, M.R. (2023). Lipid catabolism and mitochondrial uncoupling are stimulated in brown adipose tissue of amyotrophic lateral sclerosis mouse models. Genes Dis. 10, 321–324. 10.1016/j.gendis.2022.04.006.

60. Scekic-Zahirovic, J., Sendscheid, O., El Oussini, H., Jambeau, M., Sun, Y., Mersmann, S., Wagner, M., Dieterlé, S., Sinniger, J., Dirrig-Grosch, S., et al. (2016). Toxic gain of function from mutant FUS protein is crucial to trigger cell autonomous motor neuron loss. EMBO J. 35, 1077–1097. 10.15252/embj.201592559.

61. Steyn, F.J., Li, R., Kirk, S.E., Tefera, T.W., Xie, T.Y., Tracey, T.J., Kelk, D., Wimberger, E., Garton, F.C., Roberts, L., et al. (2020). Altered skeletal muscle glucose-fatty acid flux in amyotrophic lateral sclerosis. Brain Commun. 2, fcaa154. 10.1093/braincomms/fcaa154.

62. Miquel, E., Villarino, R., Martínez-Palma, L., Cassina, A., and Cassina, P. (2024). Pyruvate dehydrogenase kinase 2 knockdown restores the ability of amyotrophic lateral sclerosis-linked SOD1G93A rat astrocytes to support motor neuron survival by increasing mitochondrial respiration. Glia 72, 999–1011. 10.1002/glia.24516.

63. Li, D., Zhang, F., Zhang, X., Xue, C., Namwanje, M., Fan, L., Reilly, M.P., Hu, F., and Qiang, L. (2016). Distinct Functions of PPARγ Isoforms in Regulating Adipocyte Plasticity. Biochem. Biophys. Res. Commun. 481, 132–138. 10.1016/j.bbrc.2016.10.152.

64. Scaricamazza, S., Salvatori, I., Giacovazzo, G., Loeffler, J.P., Rene, F., Rosina, M., Quessada, C., Proietti, D., Heil, C., Rossi, S., et al. (2020). Skeletal-Muscle Metabolic Reprogramming in ALS-SOD1(G93A) Mice Predates Disease Onset and Is A Promising Therapeutic Target. iScience 23, 101087. 10.1016/j.isci.2020.101087.

65. C, P., Ml, M., G, B., R, A., C, R., Mc, S., A, B., and R, S. (2017). Absolute quantification of myosin heavy chain isoforms by selected reaction monitoring can underscore skeletal muscle changes in a mouse model of amyotrophic lateral sclerosis. Anal. Bioanal. Chem. 409. 10.1007/s00216-016-0160-2.

66. Frey, D., Schneider, C., Xu, L., Borg, J., Spooren, W., and Caroni, P. (2000). Early and selective loss of neuromuscular synapse subtypes with low sprouting competence in motoneuron diseases. J. Neurosci. Off. J. Soc. Neurosci. 20, 2534–2542. 10.1523/JNEUROSCI.20-07-02534.2000.

67. Hegedus, J., Putman, C.T., Tyreman, N., and Gordon, T. (2008). Preferential motor unit loss in the SOD1 G93A transgenic mouse model of amyotrophic lateral sclerosis. J. Physiol. 586, 3337–3351. 10.1113/jphysiol.2007.149286.

68. Lanznaster, D., Bruno, C., Bourgeais, J., Emond, P., Zemmoura, I., Lefèvre, A., Reynier, P., Eymieux, S., Blanchard, E., Vourc’h, P., et al. (2022). Metabolic Profile and Pathological Alterations in the Muscle of Patients with Early-Stage Amyotrophic Lateral Sclerosis. Biomedicines 10, 1307. 10.3390/biomedicines10061307.

69. Salvatori, I., Nesci, V., Spalloni, A., Marabitti, V., Muzzi, M., Zenuni, H., Scaricamazza, S., Rosina, M., Fenili, G., Goglia, M., et al. (2024). Trimetazidine Improves Mitochondrial Dysfunction in SOD1G93A Cellular Models of Amyotrophic Lateral Sclerosis through Autophagy Activation. Int. J. Mol. Sci. 25, 3251. 10.3390/ijms25063251.

70. Nd, P., and Bj, T. (2016). AMPK Signalling and Defective Energy Metabolism in Amyotrophic Lateral Sclerosis. Neurochem. Res. 41. 10.1007/s11064-015-1665-3.

71. S, S., I, S., S, A., V, N., A, T., G, G., A, P., M, G., N, C., L, P., et al. (2022). Repurposing of Trimetazidine for amyotrophic lateral sclerosis: A study in SOD1G93A mice. Br. J. Pharmacol. 179. 10.1111/bph.15738.

72. Pisani, D.F., Barquissau, V., Chambard, J.-C., Beuzelin, D., Ghandour, R.A., Giroud, M., Mairal, A., Pagnotta, S., Cinti, S., Langin, D., et al. (2018). Mitochondrial fission is associated with UCP1 activity in human brite/beige adipocytes. Mol. Metab. 7, 35–44. 10.1016/j.molmet.2017.11.007.

73. Wikstrom, J.D., Mahdaviani, K., Liesa, M., Sereda, S.B., Si, Y., Las, G., Twig, G., Petrovic, N., Zingaretti, C., Graham, A., et al. (2014). Hormone-induced mitochondrial fission is utilized by brown adipocytes as an amplification pathway for energy expenditure. EMBO J. 33, 418–436. 10.1002/embj.201385014.

74. Pereira, R.O., Marti, A., Olvera, A.C., Tadinada, S.M., Bjorkman, S.H., Weatherford, E.T., Morgan, D.A., Westphal, M., Patel, P.H., Kirby, A.K., et al. (2021). OPA1 deletion in brown adipose tissue improves thermoregulation and systemic metabolism via FGF21. eLife 10, e66519. 10.7554/eLife.66519.

75. Bean, C., Audano, M., Varanita, T., Favaretto, F., Medaglia, M., Gerdol, M., Pernas, L., Stasi, F., Giacomello, M., Herkenne, S., et al. (2021). The mitochondrial protein Opa1 promotes adipocyte browning that is dependent on urea cycle metabolites. Nat. Metab. 3, 1633–1647. 10.1038/s42255-021-00497-2.

76. McCluskey, G., Morrison, K.E., Donaghy, C., Rene, F., Duddy, W., and Duguez, S. (2022). Extracellular Vesicles in Amyotrophic Lateral Sclerosis. Life Basel Switz. 13, 121. 10.3390/life13010121.

77. Roy, J., Saucier, D., O’Connell, C., and Morin, P.J. (2019). Extracellular vesicles and their diagnostic potential in amyotrophic lateral sclerosis. Clin. Chim. Acta Int. J. Clin. Chem. 497, 27–34. 10.1016/j.cca.2019.07.012.

78. Wang, K., Li, Y., Ren, C., Wang, Y., He, W., and Jiang, Y. (2021). Extracellular Vesicles as Innovative Treatment Strategy for Amyotrophic Lateral Sclerosis. Front. Cell Dev. Biol. 9, 754630. 10.3389/fcell.2021.754630.

79. Banks, W.A., Sharma, P., Bullock, K.M., Hansen, K.M., Ludwig, N., and Whiteside, T.L. (2020). Transport of Extracellular Vesicles across the Blood-Brain Barrier: Brain Pharmacokinetics and Effects of Inflammation. Int. J. Mol. Sci. 21, 4407. 10.3390/ijms21124407.

80. Sykova, E., Cizkova, D., and Kubinova, S. (2021). Mesenchymal Stem Cells in Treatment of Spinal Cord Injury and Amyotrophic Lateral Sclerosis. Front. Cell Dev. Biol. 9, 695900. 10.3389/fcell.2021.695900.

81. Liu, X., Shen, L., Wan, M., Xie, H., and Wang, Z. (2024). Peripheral extracellular vesicles in neurodegeneration: pathogenic influencers and therapeutic vehicles. J. Nanobiotechnology 22, 170. 10.1186/s12951-024-02428-1.

82. Vatsa, P., Negi, R., Ansari, U.A., Khanna, V.K., and Pant, A.B. (2022). Insights of Extracellular Vesicles of Mesenchymal Stem Cells: a Prospective Cell-Free Regenerative Medicine for Neurodegenerative Disorders. Mol. Neurobiol. 59, 459–474. 10.1007/s12035-021-02603-7.

83. Katsuda, T., Oki, K., and Ochiya, T. (2015). Potential application of extracellular vesicles of human adipose tissue-derived mesenchymal stem cells in Alzheimer’s disease therapeutics. Methods Mol. Biol. Clifton NJ 1212, 171–181. 10.1007/7651_2014_98.

84. Bi, Y., Lin, X., Liang, H., Yang, D., Zhang, X., Ke, J., Xiao, J., Chen, Z., Chen, W., Zhang, X., et al. (2020). Human Adipose Tissue-Derived Mesenchymal Stem Cells in Parkinson’s Disease: Inhibition of T Helper 17 Cell Differentiation and Regulation of Immune Balance Towards a Regulatory T Cell Phenotype. Clin. Interv. Aging 15, 1383– 1391. 10.2147/CIA.S259762.

85. Garcia-Contreras, M., and Thakor, A.S. (2021). Human adipose tissue-derived mesenchymal stem cells and their extracellular vesicles modulate lipopolysaccharide activated human microglia. Cell Death Discov. 7, 98. 10.1038/s41420-021-00471-7.

86. Bonafede, R., Turano, E., Scambi, I., Busato, A., Bontempi, P., Virla, F., Schiaffino, L., Marzola, P., Bonetti, B., and Mariotti, R. (2020). ASC-Exosomes Ameliorate the Disease Progression in SOD1(G93A) Murine Model Underlining Their Potential Therapeutic Use in Human ALS. Int. J. Mol. Sci. 21. 10.3390/ijms21103651.

87. Yildirim, S., Oylumlu, E., Ozkan, A., Sinen, O., Bulbul, M., Goksu, E.T., Ertosun, M.G., and Tanriover, G. (2023). Zinc (Zn) and adipose-derived mesenchymal stem cells (AD-MSCs) on MPTP-induced Parkinson’s disease model: A comparative evaluation of behavioral and immunohistochemical results. Neurotoxicology 97, 1–11. 10.1016/j.neuro.2023.05.002.

88. Lee, M., Ban, J.-J., Kim, K.Y., Jeon, G.S., Im, W., Sung, J.-J., and Kim, M. (2016). Adipose-derived stem cell exosomes alleviate pathology of amyotrophic lateral sclerosis in vitro. Biochem. Biophys. Res. Commun. 479, 434–439. 10.1016/j.bbrc.2016.09.069.

89. Brestoff, J.R., Wilen, C.B., Moley, J.R., Li, Y., Zou, W., Malvin, N.P., Rowen, M.N., Saunders, B.T., Ma, H., Mack, M.R., et al. (2021). Intercellular Mitochondria Transfer to Macrophages Regulates White Adipose Tissue Homeostasis and Is Impaired in Obesity. Cell Metab. 33, 270–282.e8. 10.1016/j.cmet.2020.11.008.

90. Henriques, F., Bedard, A.H., Guilherme, A., Kelly, M., Chi, J., Zhang, P., Lifshitz, L.M., Bellvé, K., Rowland, L.A., Yenilmez, B., et al. (2020). Single-Cell RNA Profiling Reveals Adipocyte to Macrophage Signaling Sufficient to Enhance Thermogenesis. Cell Rep. 32, 107998. 10.1016/j.celrep.2020.107998.

91. Qiu, Y., Nguyen, K.D., Odegaard, J.I., Cui, X., Tian, X., Locksley, R.M., Palmiter, R.D., and Chawla, A. (2014). Eosinophils and Type 2 Cytokine Signaling in Macrophages Orchestrate Development of Functional Beige Fat. Cell 157, 1292–1308. 10.1016/j.cell.2014.03.066.

92. C, C., Jb, F., S, L., N, J., Cm, G., Al, G., Ya, A., Ha, S., R, G., Y, A., et al. (2021). Extracellular vesicle-based interorgan transport of mitochondria from energetically stressed adipocytes. Cell Metab. 33. 10.1016/j.cmet.2021.08.002.

93. Wilhelmsen, A., Tsintzas, K., and Jones, S.W. (2021). Recent advances and future avenues in understanding the role of adipose tissue cross talk in mediating skeletal muscle mass and function with ageing. GeroScience 43, 85–110. 10.1007/s11357-021-00322-4.

94. Kulaj, K., Harger, A., Bauer, M., Caliskan, Ö.S., Gupta, T.K., Chiang, D.M., Milbank, E., Reber, J., Karlas, A., Kotzbeck, P., et al. (2023). Adipocyte-derived extracellular vesicles increase insulin secretion through transport of insulinotropic protein cargo. Nat. Commun. 14, 709. 10.1038/s41467-023-36148-1.

95. O’Leary, M.F., Wallace, G.R., Davis, E.T., Murphy, D.P., Nicholson, T., Bennett, A.J., Tsintzas, K., and Jones, S.W. (2018). Obese subcutaneous adipose tissue impairs human myogenesis, particularly in old skeletal muscle, via resistin-mediated activation of NFκB. Sci. Rep. 8, 15360. 10.1038/s41598-018-33840-x.

96. Sheng, C.H., Du, Z.W., Song, Y., Wu, X.D., Zhang, Y.C., Wu, M., Wang, Q., and Zhang, G.Z. (2013). Human resistin inhibits myogenic differentiation and induces insulin resistance in myocytes. BioMed Res. Int. 2013, 804632. 10.1155/2013/804632.

97. Pellegrinelli, V., Rouault, C., Rodriguez-Cuenca, S., Albert, V., Edom-Vovard, F., Vidal-Puig, A., Clément, K., Butler-Browne, G.S., and Lacasa, D. (2015). Human Adipocytes Induce Inflammation and Atrophy in Muscle Cells During Obesity. Diabetes 64, 3121– 3134. 10.2337/db14-0796.

98. Wang, Y., Li, Y., Wang, X., Zhang, D., Zhang, H., Wu, Q., He, Y., Wang, J., Zhang, L., Xia, H., et al. (2013). Circulating miR-130b mediates metabolic crosstalk between fat and muscle in overweight/obesity. Diabetologia 56, 2275–2285. 10.1007/s00125-013-2996-8.

99. Yu, Y., Du, H., Wei, S., Feng, L., Li, J., Yao, F., Zhang, M., Hatch, G.M., and Chen, L. (2018). Adipocyte-Derived Exosomal MiR-27a Induces Insulin Resistance in Skeletal Muscle Through Repression of PPARγ. Theranostics 8, 2171–2188. 10.7150/thno.22565.

100. Wiedemann, F.R., Winkler, K., Kuznetsov, A.V., Bartels, C., Vielhaber, S., Feistner, H., and Kunz, W.S. (1998). Impairment of mitochondrial function in skeletal muscle of patients with amyotrophic lateral sclerosis. J. Neurol. Sci. 156, 65–72. 10.1016/s0022-510x(98)00008-2.

101. Capitanio, D., Vasso, M., Ratti, A., Grignaschi, G., Volta, M., Moriggi, M., Daleno, C., Bendotti, C., Silani, V., and Gelfi, C. (2012). Molecular signatures of amyotrophic lateral sclerosis disease progression in hind and forelimb muscles of an SOD1(G93A) mouse model. Antioxid. Redox Signal. 17, 1333–1350. 10.1089/ars.2012.4524.

102. Ghiasi, P., Hosseinkhani, S., Noori, A., Nafissi, S., and Khajeh, K. (2012). Mitochondrial complex I deficiency and ATP/ADP ratio in lymphocytes of amyotrophic lateral sclerosis patients. Neurol. Res. 34, 297–303. 10.1179/1743132812Y.0000000012.

103. Salvatori, I., Ferri, A., Scaricamazza, S., Giovannelli, I., Serrano, A., Rossi, S., D’Ambrosi, N., Cozzolino, M., Giulio, A.D., Moreno, S., et al. (2018). Differential toxicity of TAR DNA-binding protein 43 isoforms depends on their submitochondrial localization in neuronal cells. J. Neurochem. 146, 585–597. 10.1111/jnc.14465.

104. Wang, W., Wang, L., Lu, J., Siedlak, S.L., Fujioka, H., Liang, J., Jiang, S., Ma, X., Jiang, Z., da Rocha, E.L., et al. (2016). The inhibition of TDP-43 mitochondrial localization blocks its neuronal toxicity. Nat. Med. 22, 869–878. 10.1038/nm.4130.

105. Remels, A.H.V., Langen, R.C.J., Schrauwen, P., Schaart, G., Schols, A.M.W.J., and Gosker, H.R. (2010). Regulation of mitochondrial biogenesis during myogenesis. Mol. Cell. Endocrinol. 315, 113–120. 10.1016/j.mce.2009.09.029.

106. Rahman, F.A., and Quadrilatero, J. (2021). Mitochondrial network remodeling: an important feature of myogenesis and skeletal muscle regeneration. Cell. Mol. Life Sci. 78, 4653–4675. 10.1007/s00018-021-03807-9.

107. Motohashi, N., Minegishi, K., and Aoki, Y. (2023). Inherited myogenic abilities in muscle precursor cells defined by the mitochondrial complex I-encoding protein. Cell Death Dis. 14, 1–13. 10.1038/s41419-023-06192-2.

108. Chabi, B., Hennani, H., Cortade, F., and Wrutniak-Cabello, C. (2021). Characterization of mitochondrial respiratory complexes involved in the regulation of myoblast differentiation. Cell Biol. Int. 45, 1676–1684. 10.1002/cbin.11602.

109. Kong, X., Yao, T., Zhou, P., Kazak, L., Tenen, D., Lyubetskaya, A., Dawes, B.A., Tsai, L., Kahn, B.B., Spiegelman, B.M., et al. (2018). Brown Adipose Tissue Controls Skeletal Muscle Function via the Secretion of Myostatin. Cell Metab. 28, 631–643.e3. 10.1016/j.cmet.2018.07.004.

110. Wang, L., Zheng, Z.-G., Meng, L., Zhu, L., Li, P., Chen, J., and Yang, H. (2021). Statins induce skeletal muscle atrophy via GGPP depletion-dependent myostatin overexpression in skeletal muscle and brown adipose tissue. Cell Biol. Toxicol. 37, 441–460. 10.1007/s10565-020-09558-w.

111. Rome, S. (2022). Muscle and Adipose Tissue Communicate with Extracellular Vesicles. Int. J. Mol. Sci. 23, 7052. 10.3390/ijms23137052.

112. Figliolini, F., Ranghino, A., Grange, C., Cedrino, M., Tapparo, M., Cavallari, C., Rossi, A., Togliatto, G., Femminò, S., Gugliuzza, M.V., et al. (2020). Extracellular Vesicles From Adipose Stem Cells Prevent Muscle Damage and Inflammation in a Mouse Model of Hind Limb Ischemia. Arterioscler. Thromb. Vasc. Biol. 40, 239–254. 10.1161/ATVBAHA.119.313506.

113. Perez-Riverol, Y., Bai, J., Bandla, C., García-Seisdedos, D., Hewapathirana, S., Kamatchinathan, S., Kundu, D.J., Prakash, A., Frericks-Zipper, A., Eisenacher, M., et al. (2022). The PRIDE database resources in 2022: a hub for mass spectrometry-based proteomics evidences. Nucleic Acids Res. 50, D543–D552. 10.1093/nar/gkab1038.

